# Structures of trehalose-6-phosphate synthase, Tps1, from the fungal pathogen *Cryptococcus neoformans*: a target for novel antifungals

**DOI:** 10.1101/2023.03.14.530545

**Authors:** Erica J. Washington, Ye Zhou, Allen L. Hsu, Matthew Petrovich, Jennifer L. Tenor, Dena L. Toffaletti, Ziqiang Guan, John R. Perfect, Mario J. Borgnia, Alberto Bartesaghi, Richard G. Brennan

## Abstract

Invasive fungal diseases are a major threat to human health, resulting in more than 1.5 million annual deaths worldwide. The arsenal of antifungal therapeutics remains limited and is in dire need of novel drugs that target additional biosynthetic pathways that are absent from humans. One such pathway involves the biosynthesis of trehalose. Trehalose is a disaccharide that is required for pathogenic fungi to survive in their human hosts. In the first step of trehalose biosynthesis, trehalose-6-phosphate synthase (Tps1) converts UDP-glucose and glucose-6-phosphate to trehalose-6-phosphate. Here, we report the structures of full-length *Cryptococcus neoformans* Tps1 (CnTps1) in unliganded form and in complex with uridine diphosphate and glucose-6-phosphate. Comparison of these two structures reveals significant movement towards the catalytic pocket by the N-terminus upon ligand binding and identifies residues required for substrate-binding, as well as residues that stabilize the tetramer. Intriguingly, an intrinsically disordered domain (IDD), which is conserved amongst Cryptococcal species and closely related Basidiomycetes, extends from each subunit of the tetramer into the “solvent” but is not visible in density maps. We determined that the IDD is not required for *C. neoformans* Tps1-dependent thermotolerance and osmotic stress survival. Studies with UDP-galactose highlight the exquisite substrate specificity of CnTps1. *In toto*, these studies expand our knowledge of trehalose biosynthesis in *Cryptococcus* and highlight the potential of developing antifungal therapeutics that disrupt the synthesis of this disaccharide or the formation of a functional tetramer and the use of cryo-EM in the structural characterization of CnTps1-ligand/drug complexes.

**Significance Statement:** Fungal infections are responsible for over a million deaths worldwide each year. Biosynthesis of a disaccharide, trehalose, is required for multiple pathogenic fungi to transition from the environment to the human host. Enzymes in the trehalose biosynthesis pathway are absent in humans and, therefore, are potentially significant targets for novel antifungal therapeutics. One enzyme in the trehalose biosynthesis is trehalose-6-phosphate synthase (Tps1). Here, we describe the cryo-electron microscopy structures of the CnTps1 homo-tetramer in the unliganded form and in complex with a substrate and a product. These structures and subsequent biochemical analysis reveal key details of substrate-binding residues and substrate specificity. These structures should facilitate structure-guided design of inhibitors against CnTps1.

## Introduction

Invasive fungal diseases (IFDs) caused by pathogenic fungi such as *Cryptococcus*, *Candida* and *Aspergillus* are a major threat to human health, resulting in more than one and a half million deaths worldwide each year (1, 2). These mortality rates are surprisingly high, given the fact that fungal infections are usually associated with superficial infections of skin and nails. However, IFDs result in increased mortality in immunocompromised populations, including HIV-infected patients and patients receiving immunosuppressive therapies (1–3). Additionally, the severe acute respiratory syndrome coronavirus 2 (SARS-CoV-2; COVID-19) pandemic has resulted in a significant population of COVID-19 positive patients with increased susceptibility to IFDs, as has been demonstrated for pulmonary aspergillus and invasive candidiasis infections (4–10).

Unfortunately, the current arsenal of antifungal drugs, consisting of the polyenes, azoles and echinocandins, is insufficient to manage the mortality caused by IFDs due to significant off-target effects, the rapid emergence of resistance to antifungal therapeutics and the emergence of intrinsically drug resistant fungal pathogens, such as *Candida auris* and *Candida glabrata* (11–14) Therefore, in addition to the development of new formulations of current antifungal drugs, there is a critical need for the development of new classes of broad-spectrum antifungal drugs that are fast-acting and safe. The identification of targets that are not present in the human host is critical to the development of antifungal drugs with low toxicity. This is often difficult, as both fungi and humans are eukaryotes. However, pathways involved in the fungal stress response have been implicated recently as key targets, including the trehalose biosynthesis pathway (14–17).

Trehalose is a non-reducing disaccharide composed of two glucose molecules linked by an α,α-1,1-glycosidic bond. Fungal cells synthesize trehalose to protect their proteins and membranes from stresses encountered during the infection process (18–25). The canonical trehalose biosynthesis machinery in pathogenic fungi commences with trehalose-6-phosphate synthase, Tps1, a glucosyltransferase that converts uridine diphosphate glucose (UDPG) and glucose-6-phosphate (G6P) to trehalose-6-phosphate (T6P). Subsequently, trehalose-6-phosphate phosphatase (Tps2), removes the phosphate group to generate the final product of trehalose (Figure 1A). In *Candida albicans* and *Aspergillus fumigatus* there is an additional protein in the pathway, Tps3, with no known enzymatic function. The machinery to synthesize trehalose is found in plants, insects, fungi and bacteria, but not in humans (26). For this reason, an antifungal therapeutic that targets the trehalose biosynthesis pathway could well result in a drug with minimal off-target effects, leading to greatly reduced toxicity in patients. Indeed, disruption of the *TPS1* gene in *Cryptococcus neoformans* results in fungi that are avirulent in mice and rabbits and, interestingly, disruption of *TPS2* was followed by the accumulation of the toxic intermediate trehalose 6-phosphate, causing fungal cell death (27). Additionally, in *C. neoformans* trehalose acts as a stress protectant and is required for growth at high temperatures (27). Similarly, disruption of the *TPS1* and *TPS2* genes in *Cryptococcus gattii* results in decreased infectivity in mice (28). Similar phenotypes indicating the importance of Tps1 and Tps2 to cell survival and sometimes virulence are also seen in trehalose biosynthesis mutants in *C. albicans* (29–31) and *A. fumigatus* (32, 33). Therefore, we hypothesize that targeting the trehalose biosynthesis pathway will result in the development of a novel broader-spectrum antifungal drug, which would be both highly effective and, importantly, safe for use in immunocompromised patients.

**Figure 1.**
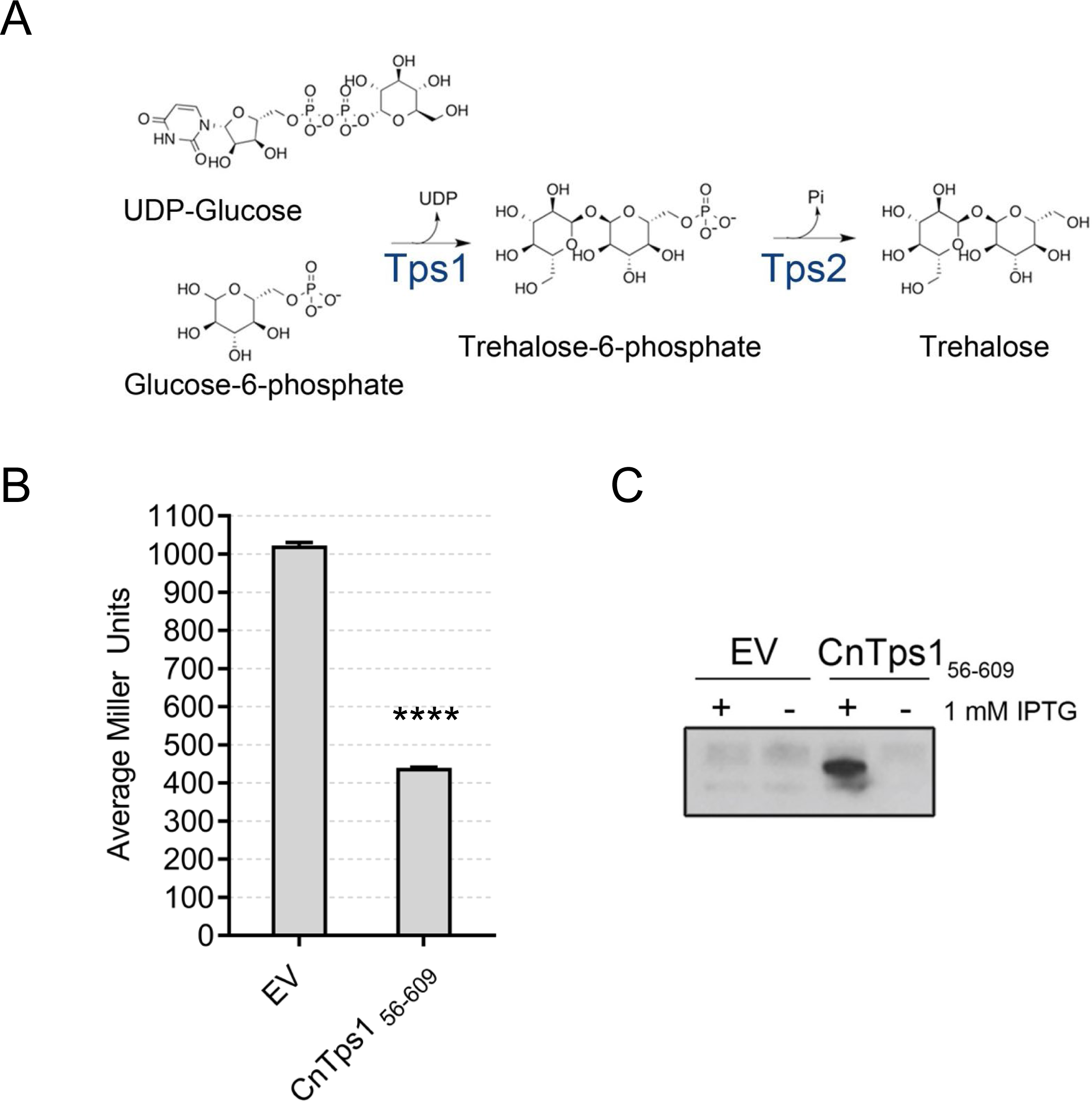
Trehalose-6-phosphate synthase (Tps1) from *Cryptococcus neoformans* self-associates. **A)** Schematic of the canonical trehalose biosynthesis pathway in fungi. **B)** Bacterial two-hybrid results indicate a self-association of LexA-CnTps1_56-669_ by a reduction in β-galactoside activity after induction of expression with 1 mM IPTG. Error bars represent standard error of triplicate biological replicates (N=3). Statistically significant differences were demonstrated with an unpaired Student’s *t*-test (**** *P* < 0.0001). **C)** Anti-LexA Western blot confirms the expression of LexA-CnTps1_56-669_.

The trehalose biosynthesis pathway and its contribution to the response of fungi to the increasingly warmer global temperatures is also important to understand. Climate change plays a critical role in the emergence of fungal diseases by reducing the threshold between environmental temperatures and mammalian basal temperatures (34–38). When fungi, especially those possessing virulence factors, acquire thermotolerance they also acquire the possibility of becoming more pathogenic. Importantly, the trehalose biosynthesis pathway in *C. neoformans* plays a role in thermotolerance (39). CnTps1 is required for *C. neoformans* to grow at higher temperatures (28). Similarly, the ability of trehalose to accumulate in *C. neoformans* grown at 37 °C is dependent on CnTps1 (28). Therefore, understanding the function of CnTps1 and its structure should help us understand how environmental fungi evolve thermotolerance to become pathogenic.

Pertinent to the development of an antifungal therapeutic which targets the trehalose biosynthesis pathway is an intimate knowledge and understanding of the structures of enzymes and proteins involved in this critical pathway. Several structures of Tps1 and Tps2 from nematodes, bacteria and pathogenic fungi have been determined using x-ray crystallography (40–49). Subsequently, compounds that inhibit trehalose biosynthesis enzymes functions by targeting their catalytic pockets have been investigated but not yet advanced for clinical use (45, 50–55). As a key starting point to develop inhibitors of CnTps1, we describe structures of this 669-residue (74 kDa per protomer) enzyme in both its unliganded (substrate-free) form and in complex with UDP and G6P using single particle cryo-electron microscopy (cryo-EM). Unlike Tps1 from *Candida albicans* (46), CnTps1 has three major insertions, 55 residues at the N-terminus, 63 residues at the C-terminus and a 92-residue intrinsically disordered domain (IDD) inserted within the core of the enzyme, all of unknown function. Residues located in the catalytic pocket and the IDD were assayed for their contribution to the activity of this enzyme. Substrate specificity was investigated further and revealed the exquisite ability of CnTps1 to discriminate at the level of a single interaction. Hence, this work lays the groundwork for subsequent CnTps1 structure-based drug design.

## Results

### Trehalose biosynthesis protein CnTps1 self-associates in bacterial two-hybrid assays

The trehalose biosynthesis pathway is a two-step process. Tps1, trehalose-6-phosphate synthase, converts uridine diphosphate glucose (UDPG) and glucose-6-phosphate (G6P) into trehalose-6-phosphate (T6P). The phosphate group is subsequently removed from T6P by trehalose-6-phosphate phosphatase (Tps2) to yield trehalose (Figure 1A) (17). Previously the structures of Tps1/OtsA from *Candida albicans* (46), *Aspergillus fumigatus* (46), *E. coli* (*41, 43, 44, 49*), *Streptomyces venezuelae* (40) and *Magnaporthe oryzae* (49) were determined using x-ray crystallography. Additionally, the structure of the catalytically inactive Tps1-like N-terminal domain of *C. albicans* Tps2 was determined (47).

Inspection of the Tps1 crystal structures from *Candida albicans* (46), *Aspergillus fumigatus* (46), *E. coli* (43, 44), *Streptomyces venezuelae* (40) and *Magnaporthe oryzae* (49) and the catalytically inactive Tps1-like N-terminal domain of *C. albicans* Tps2 (47) revealed that Tps1 most likely exists as a homo-tetramer. To determine if subunits of the *Cryptococcus neoformans* var. *grubii* strain H99 Tps1, hereafter referred to as CnTps1, self-associate, a bacterial two-hybrid system based on the LexA DNA-binding repressor was established (56). Expression of full-length LexA-CnTps1 in the bacterial two-hybrid system was not detected. However, LexA-CnTps1_56-669_, in which the unstructured 55 N-terminal residues of CnTps1 were deleted, does self-associate in the bacterial two-hybrid assay, as evidenced by a reduction in β-galactosidase activity after protein expression was induced with 1 mM IPTG (Figure 1B). An anti-LexA Western blot shows the expression of LexA-CnTps1_56-669_ at the appropriate molecular weight (Figure 1C). These data indicate that CnTps1 can self-associate at physiological concentrations and support the hypothesis that CnTps1 forms homo-oligomers.

### The structure of the unliganded (substrate-free) CnTps1 homo-tetramer

CnTps1 is approximately 200 residues larger than other Tps1 orthologues, due to the addition of a 55-residue N-terminal extension, a 63-residue C-terminal extension and an internal insertion from residues M209 to I300 (Supplemental Figure 1). Each extension is predicted to be structurally disordered. In order to determine the structure of CnTps1, the full-length (74 kDa, 669-residue per protomer) CnTps1 protein was expressed with an N-terminal hexahistidine tag and purified from *E. coli* via Ni^2+^-NTA affinity chromatography (Supplemental Figure 2A). Following confirmation that 6xHis-CnTps1 was enzymatically active (Supplemental Figure 2B), the protein sample was subjected to single particle cryo-electron microscopy analysis. 2,520 micrographs were collected from grids prepared with 6xHis-CnTps1 (Supplemental Figure 3).

No additional ligands were added to the protein sample. 2D classification, performed on 907 micrographs with resolution higher than 4.0 Å, revealed tetrameric CnTps1 particles (Supplemental Figure 3). 3D classification and initial refinement steps were performed without imposing symmetry (Supplemental Figure 3), resulting in a 5.3 Å resolution map. Additional refinement and polishing steps, including adding D2 symmetry, resulted in an unliganded CnTps1 map at 3.3 Å global resolution, as estimated by the “gold standard” Fourier shell correlation (FSC) = 0.143 criteria (Data Table 1 and Supplemental Figure 4A,B). Four protomers can be readily observed in the unliganded CnTps1 cryo-EM map (Supplemental Figure 4C). The final map shows features of well-resolved side chains of many amino acid residues (Supplemental Figure 4D). The atomic model for unliganded CnTps1, which was built based on the cryo-EM reconstruction, supports the conclusion that unliganded CnTps1 forms a homo-tetramer (Figure 2).

**Figure 2.**
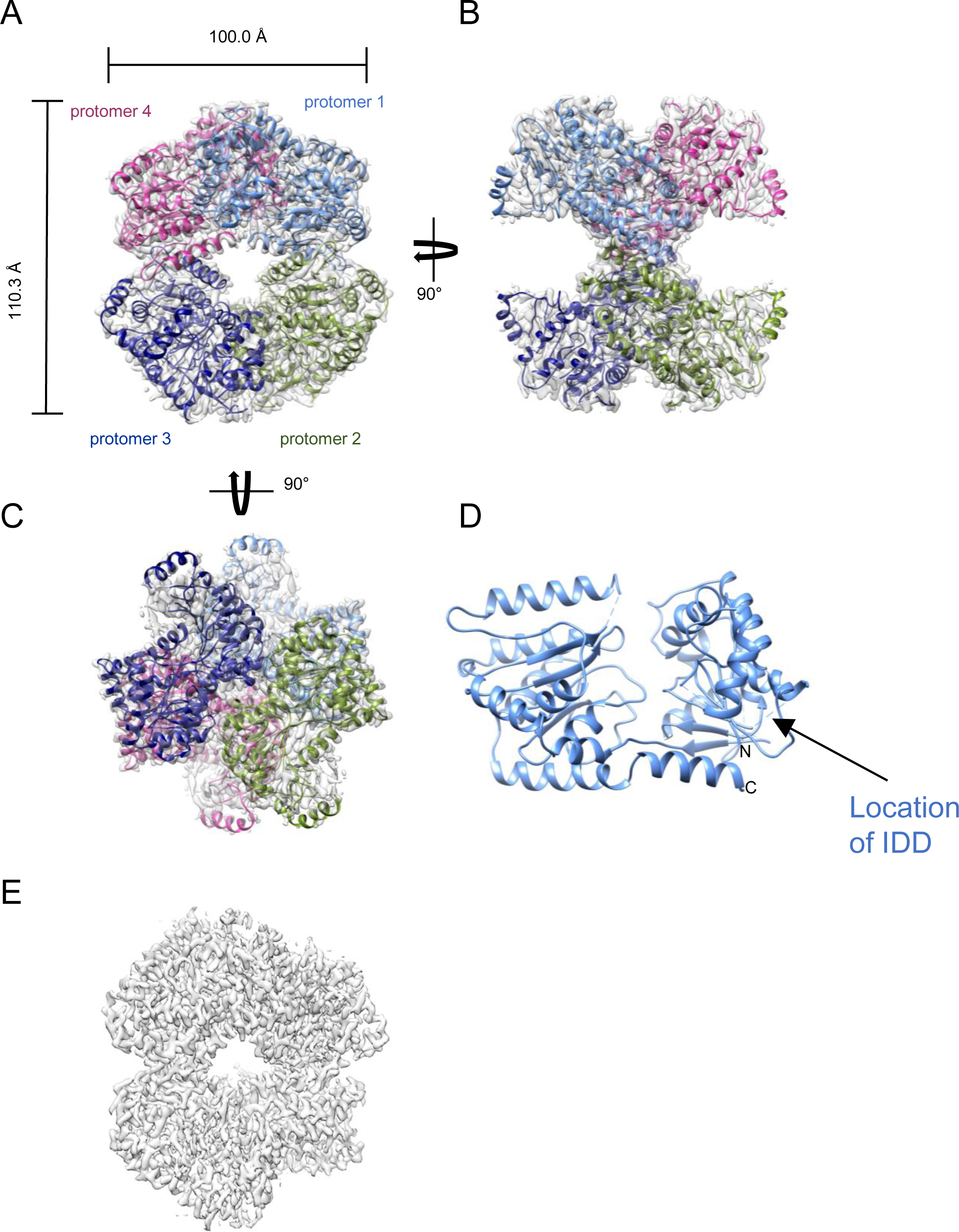
Structure of the unliganded *Cryptococcus neoformans* Tps1 homo-tetramer. **A)** Structure of the unliganded *C. neoformans* Tps1 homo-tetramer. The density map is shown in grey, overlayed with the model as a ribbon diagram. The protomers are colored and labelled in light blue, green, navy and magenta. The dimensions of the “front” view of the tetramer are labelled in Å. **B)** Structure of the tetramer viewed after a 90° rotation around the vertical axis. **C)** Structure of the tetramer viewed after a 90° rotation around the horizontal axis. **D)** Ribbon diagram of a single protomer of the unliganded CnTps1 cryo-EM structure with the N and C-termini labelled. **E)** Density for unliganded CnTps1 is shown in grey.

The four CnTps1 protomers, designated as protomers 1 – 4, assemble into the tetrameric structure with dimensions of 100.0 x 110.3 x 107.6 Å (Figure 2A-C). The four protomers interact via two α-helical domains that are found in the central region of the homo-tetramer (Figure 2A). The unbound substrate-binding pockets, are open and outward-facing, and therefore accessible to ligands and substrates (Figure 2A,B). The “bottom-facing” view of the homo-tetramer shows a staggered appearance of the top and bottom pair of protomers (Figure 2C).

Each CnTps1 protomer is comprised of N and C-terminal lobes, containing residues 1-378 and 379-669, respectively. Both lobes are spanned by an α-helix, formed by residues 572-603 (Figure 2D). Each domain contains a modified Rossman fold that is characteristic of the retaining glycosyltransferase family (46, 57) (Figure 2D) and similar to the published CaTps1 and CaTps1-like N-terminal domain of CaTps2 structures (Supplemental Figure 5) (40, 47, 49). A common feature that is observed in these substrate-free structures is a lack of density in portions of the N-terminus, suggestive of conformational flexibility in the absence of ligands. Indeed, the CnTps1 cryo-EM map shows clear density for secondary structures in the C-terminus, whilst the density was poor in several parts of the N-terminal domain, including that for residues 1-58, 68-100, 127, 180, 210-301, 354-357 and 364-368 (Figure 2D). Not unexpectedly, the N-terminus extension, the C-terminus and the internal insertion, residues 209-300, are not visible in this structure suggesting multiple accessible conformations or intrinsic disorder.

### The structure of the CnTps1-UDP-G6P complex

In order to identify residues in the catalytic pocket of CnTps1 as well as to visualize any conformational changes necessary for substrate binding and catalysis, we determined the cryo-EM structure of the *C. neoformans* H99 Tps1 in complex with the substrate, G6P, and a product, UDP, to 3.1 Å global resolution (Figure 3A-D). 3,182 movies were collected from grids prepared with 6xHis-CnTps1 and an excess of UDP and G6P (Supplemental Figure 6). The 2D classification revealed a tetrameric structure like those observed for unliganded CnTps1 (Supplemental Figure 6). Final refinement was performed with D2 symmetry imposed (Supplemental Figure 6). The addition of G6P and UDP resulted in a higher global resolution of the CnTps1 structure when compared to the unliganded structure (Data Table 2). The substrate-binding pocket is more ordered (Supplemental Figure 7A,B). Four protomers can be detected in the CnTps1-UDP-G6P cryo-EM map (Supplemental Figure 7C) and the final map shows features of well-resolved side chains (Supplemental Figure 7D).

**Figure 3.**
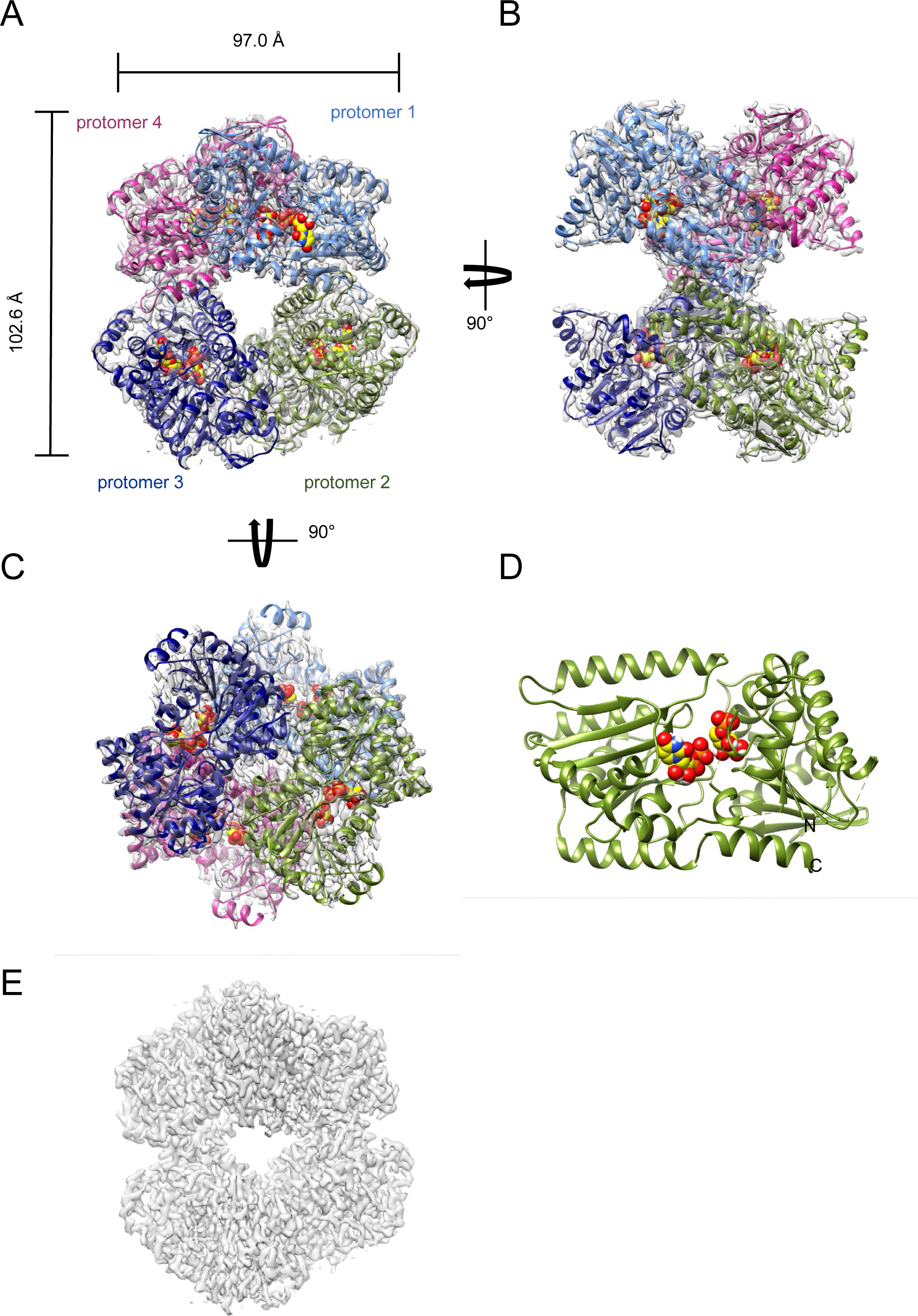
Structure of the *Cryptococcus neoformans* Tps1 homo-tetramer bound to UDP and G6P. **A)** Structure of the *C. neoformans* Tps1 homo-tetramer bound to UDP and G6P. The density map is shown in grey, overlayed with the model as a ribbon diagram. The protomers are colored and labelled in light blue, green, navy and magenta. The dimensions of the “front” view of the tetramer are labelled in Å. Ligands UDP and G6P are shown with a space-filling representation. **B)** Structure of the tetramer bound to UDP and G6P viewed after a 90° rotation around the vertical axis. **C)** Structure of the tetramer bound to UDP and G6P viewed after a 90° rotation around the horizontal axis. **D)** Ribbon diagram of a single protomer of the CnTps1-UDP-G6P structure with the N and C-termini labelled and ligands in the substrate-binding pocket shown as atom-colored, space-filling molecules. **E)** Density for CnTps1-UDP-G6P is shown in grey.

As observed in the unliganded structure, the N and C-termini and the internal insertion remain unstructured in the presence of ligands. The structure of CnTps1 bound to UDP and G6P is similar to that of the published CaTps1-UDP-G6P complex and the N-terminus of CnTps2 with root mean square deviations (rmsds) of 1.2 Å for 450 corresponding C_α_ atoms and 1.5 Å for 468 corresponding C_α_ atoms, respectively (46, 47, 58). Views of the CnTps1-UDP-G6P tetramer show the central binding region of UDP and G6P inside each protomer (Figure 3A-D). The CnTps1-UDP-G6P complex UDP bound to the C-terminal domain and the G6P adjacent to the N-terminal domain (Figure 3D). The binding sites of the ligands are in excellent agreement with the published structure of CaTps1-UDP-G6P and CaTps1-UDPG (46).

### Ligand-induced conformational changes in *C. neoformans* Tps1

Significant movement and conformational changes of the core of the N-terminal domain are observed due to ligand binding to CnTps1 (Figure 4A,B). Structural alignments of the individual N-terminal domain (residues 56 – 378) and C-terminal domain (residues 379 – 603) of the unliganded Tps1 protein onto the corresponding domains of UDP-G6P bound CnTps1 reveal rmsds of 1.1 and 0.4 Å, respectively. Superposition of the complete structure of unliganded CnTps1 onto CnTps1-UDP-G6P reveals an rmsd of 1.4 Å for 433 corresponding C_α_ atoms. These rmsd values are a result of ligand-induced conformational “closing” of CnTps1, with the movement occurring primarily in the N-terminus. Thus, unliganded CnTps1 protomers are in a more “open” conformation as compared to the ligand-bound subunits. Systematic analysis of the domain movements using the program DynDom (59, 60) shows that the large movements of the N-terminal G6P-binding lobe centered around α-helix 2 (residues 112 to 125), which is located on the surface of the N-terminal domain, result in a rotation of approximately 20° (Supplemental Figure 8). This rotation causes an inward, approximately 7 Å movement of α-helix 2 (Figure 4B). The linker that facilitates this movement includes residues 113 through 145. This rotation results in a 29% closure of the N and C-terminal lobes around the substrate-binding pocket. Interestingly, the local resolution proximal to CnTps1 α-helix 2 is the lowest in the unliganded structure, which indicates flexibility and a propensity for movement, consistent with the results of the DynDom analysis. Since the CnTps1-UDP-G6P complex contains both a product (UDP) and a substrate (G6P) as ligands, the completely closed form of the protein is likely not visualized in this structure. Regardless, the movement of this domain would facilitate the S_N_-i catalytic mechanism proposed for Tps1 proteins in the GT-B fold retaining glycosyltransferase family, including *C. albicans* Tps1 and *E. coli* OtsA (41, 46, 57, 61). Indeed, superposition of UDP onto the UDPG from the CaTps1-UDPG complex (PDB ID 5HUT) reveals that the G6P substrate bound to CnTps1 is proximal to and properly aligned with the glucose of the UDPG to allow the ready formation of the 1,1 glycosidic bond, required for the formation of T6P (Supplemental Figure 9).

**Figure 4.**
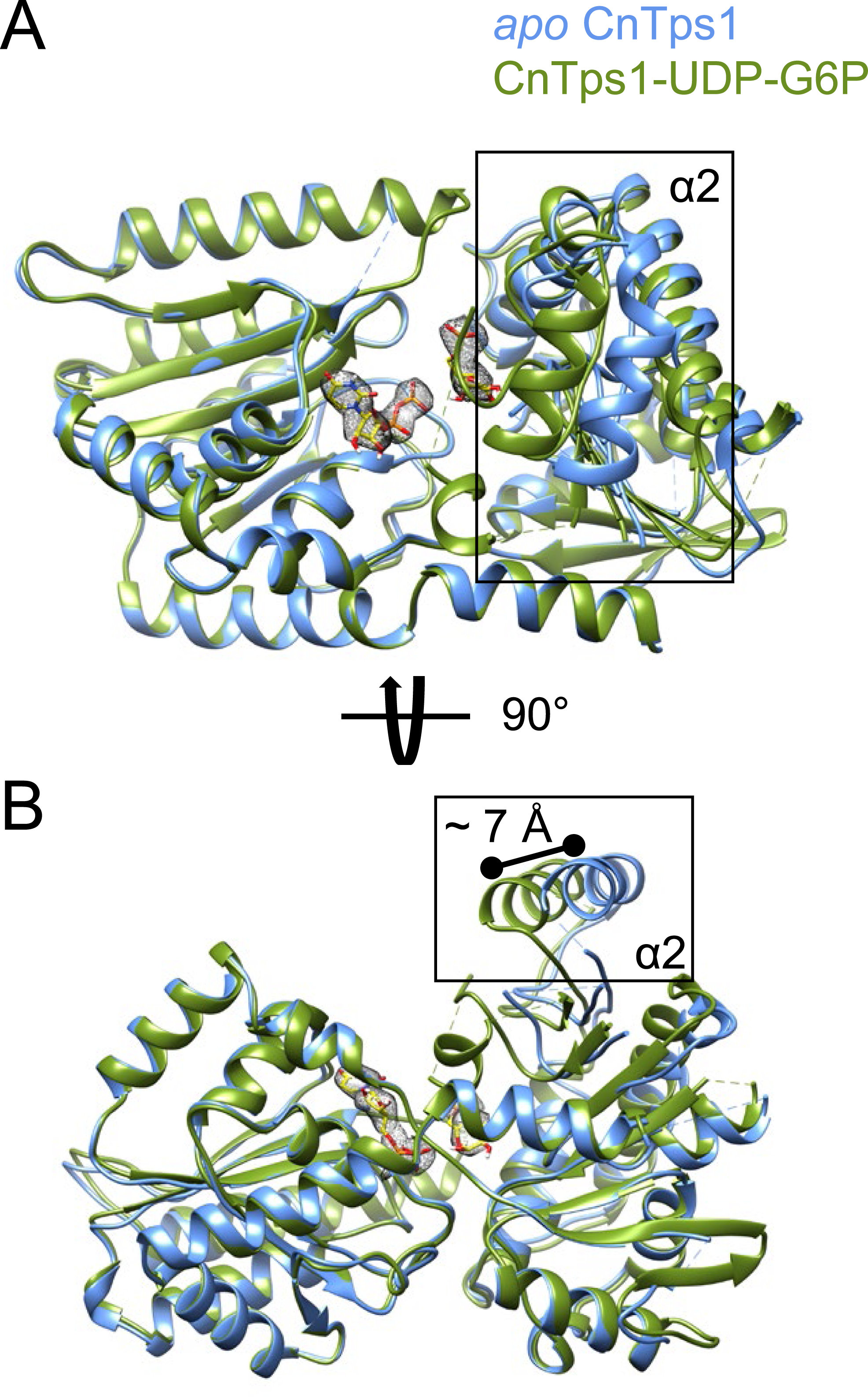
The binding of UDP and G6P induces a conformational change in CnTps1. **A)** Overlay of the unliganded CnTps1 protomer (light blue) with the CnTps1 protomer bound to UDP and G6P (green). Density (grey mesh) is shown around ligands UDP and G6P in the catalytic pocket. Position of α-helix 2 is indicated with a black box. **B)** Overlay of the unliganded CnTps1 protomer (light blue) with CnTps1-UDP-G6P (green) viewed after a 90° rotation around the horizontal axis. Electron microscopy density is shown for UDP and G6P in the substrate-binding pocket. The position of α-helix 2 is boxed, with the black line demonstrating the movement of approximately 7 Å.

### CnTps1 substrate-binding residues

We were able to detect strong density for the bound ligand and substrate-binding residues of CnTps1 (Figure 5A). The UDP molecule is bound to the C-terminal portion of each protomer (Figure 5B). The orientation of the uracil base and CnTps1 is mediated by a hydrogen bond between the exocyclic O4 and the V492 amide NH group. The O2 and O3 hydroxyl groups of the ribose ring of UDP form hydrogen bonds with the side chain of E522. The orientation of the phosphate groups of UDP in CnTps1 is mediated by several interactions. One hydrogen bond is made between the α-phosphate group and the backbone amide group of L518. In CaTps1 (46) there is a conserved Arg-Lys pair (residues R280 and K285) that interacts with the phosphates. We were unable to detect side chain density for the equivalent CnTps1 Arg, indicating movement of the substrate-binding pocket, potentially ready to release the product UDP. There is, however, a hydrogen bond between K420 and the β-phosphate. Based on an overlay with the CaTps1-UDPG structure, we predicted that the glucose moiety in the native substrate, UDPG, would be stabilized by CnTps1 residue D514 and positioned to deprotonate G6P (Figure 5B and Supplemental Figure 10A) (46). The G6P molecule is located adjacent to the N-terminal lobe in each protomer. A key interaction stabilizing G6P is residue R453, which interacts with the G6P phosphate group (Figure 5C).

**Figure 5.**
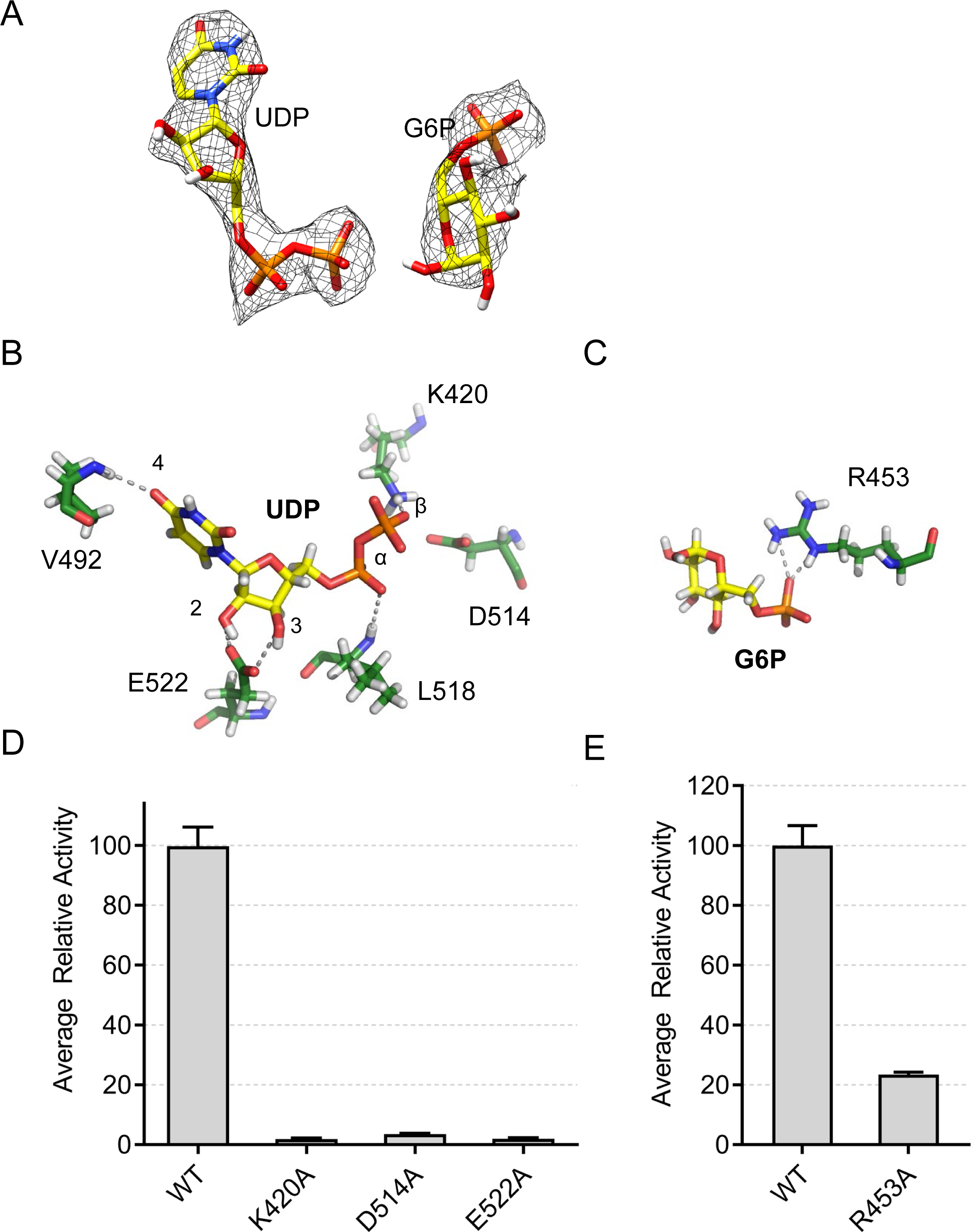
Substrate-binding residues of CnTps1. **A)** Cryo-EM density for UDP and G6P, shown as light grey mesh. UDP and G6P are shown as atom-colored sticks. **B)** View of the CnTps1 residues involved in binding UDP. UDP and residues of the C-terminal domain are shown as atom-colored sticks. Hydrogen bonds are shown by dashes. Key atoms of the UDP molecule are labelled. **C)** View of CnTps1 residue R453, which binds G6P. G6P and CnTps1 R453 are shown as atom-colored sticks. Hydrogen bonds are shown by dashes. **D)** Relative activity of wild-type CnTps1 and UDP-binding mutants. Error bars represent the standard error of three independent measurements. **E)** Relative activity of wild-type CnTps1 and the G6P-binding mutant R453A. Error bars represent the standard error of three independent measurements.

CnTps1 substrate-binding residues K420, D514, L518, E522 and R453 are conserved amongst Tps1 proteins from *C. neoformans*, *C. albicans*, *A. fumigatus*, *M. oryzae*, *S. venezuelae* and *E. coli* (Supplemental Figure 1 and Supplemental Figure 10B). In *C. albicans* Tps1 these residues have been demonstrated to be required for hyphal development and thermotolerance (46). Site-directed mutagenesis was used to generate variants of CnTps1 in which potential substrate-binding residues were mutated as follows: E522A, K420A, D514A and R453A. All CnTps1 variants eluted similarly to wild-type CnTps1 on the size exclusion column, indicating that they are properly folded (Supplemental Figure 11). Tps1 coupled enzyme assays revealed that mutation of the residues that interact with UDP/UDPG resulted in severe loss-of-function (Figure 5D). Mutation of R453 to an alanine, which is the single identified G6P-interacting residue, resulted in greatly reduced CnTps1 activity as well (Figure 5E).

### Interactions within the CnTps1 homo-tetramer

The interactions between several side chains in the interfaces between CnTps1 protomers explain the formation and stabilization of the CnTps1 homo-tetramer. The two key interfaces responsible for formation of the CnTps1 homo-tetramer are referred to as interface #1 and interface #2 (Supplemental Figure 12A). PDBePISA (62), revealed the buried surface areas for each interface are similar with 1062.9 Å buried in interface #1 and 986.2 Å buried in interface #2. The calculated free energy (ΔG) of the two interfaces is −10.7 kcal/mol and 0.1 kcal/mol, respectively, which suggests the former interface is a critical component of oligomerization, whilst the latter might allow the formation of a Tps1-Tps2 hetero-oligomer.

The key interactions in interface #1 are the hydrogen bonds between N467 and E468, as well as R403 and E481 (Supplemental Figure 12B,C). Interface #2 also has hydrogen bonds that help stabilize the homo-tetramer, including hydrogen bonds between the E314 backbone oxygen and R513 side chain (Supplemental Figure 12D). Ionic interactions between R513 and E314 and between R322 and E538 also contribute to the formation of interface #2. None of the interface residues directly contact UDP or G6P, indicating that any effect mutation of these residues might have on activity would be allosteric. Our cryo-EM results confirm that the homo-tetrameric form of CnTps1 accurately represents the tetrameric structure in solution.

### CnTps1 contains a conserved intrinsically disordered domain

CnTps1 contains three unstructured insertions (Supplemental Figure 13A). These regions have prevented the determination of its structure by x-ray crystallography, as these regions are the main differences between CnTps1 and other crystallized Tps1 orthologues. The most prominent unstructured region in CnTps1 is an internal insertion of 92 residues (residues M209-I300), referred to hereafter as the Intrinsically <u>D</u>isordered <u>D</u>omain or IDD (Supplemental Figure 13A). Interestingly, the CnTps1 IDD is found only in the Tps1 proteins in fungi within the fungal division, Basidiomycota (Supplemental Figure 13B). More specifically, the CnTps1 IDD is highly conserved within Cryptococcal species and closely related Basidiomycetes, with *C. neoformans*, *C. gattii* and *C. deneoformans* containing IDDs with greater than 92% identity (Supplemental Figure 13A,B). The CnTps1 IDD is not present in the Ascomycota division or the Mucoromycota phylum (Supplemental Figure 13B).

As anticipated, density for the CnTps1 IDD was not detected in the structures of unliganded CnTps1 or CnTps1-UDP-G6P, consistent with the idea that the IDD is conformationally flexible. However, based on the CnTps1 structures, we can determine that the IDD exits and returns to the structured regions in adjacent parts of the N-terminus of CnTps1. To determine if the IDD is required for CnTps1 activity *in vitro*, we designed and purified a chimeric protein in which the 92-residue CnTps1 IDD was replaced with the six residues (GNKKKN) of *C. albicans* Tps1 that connect α-helix 5 to β-strand 4 (Supplemental Figure 14). We refer to this protein as CnTps1 ΔIDD.

To determine the effect that the CnTps1 IDD has on CnTps1 function, we purified CnTps1 ΔIDD and confirmed that it was reduced in size compared to wild-type CnTps1, as determined by size exclusion chromatography (Supplemental Figure 11). Notably, deletion of the CnTps1 IDD did not affect formation of the tetramer (Supplemental Figure 11). Circular dichroism experiments also confirmed that CnTps1 ΔIDD is properly folded (Supplemental Figure 15). Coupled enzyme activity assays with CnTps1 ΔIDD were performed and revealed that the CnTps1 IDD is not required for enzymatic activity of CnTps1 *in vitro* (Supplemental Figure 13C). Indeed, wild-type CnTps1 and the CnTps1 ΔIDD variant have similar catalytic efficiencies (Supplemental Table 1).

CnTps1 is required for the growth of *C. neoformans* at 37 °C (27). To determine if the CnTps1 IDD is required for the CnTps1-mediated thermotolerance, strains of *C. neoformans* H99 lacking the CnTps1 IDD were generated. Wild-type *C. neoformans*, the *tps1*Δ strain and the *TPS1^IDD^*^Δ^ strain were able to grow similarly at 30 °C (Supplemental Figure 16). At 37 °C, the *C. neoformans TPS1^IDD^*^Δ^ strain had wild-type levels of growth, compared to the temperature-sensitive phenotype observed for *C. neoformans tps1*Δ (Supplemental Figure 16). Furthermore, only the *C. neoformans tps1*Δ strain could not grow at 39 °C (Supplemental Figure 16). The complementation strain of *C. neoformans TPS1^IDD^*^Δ^ strain expressing the wild-type *TPS1* grew like wild-type *C. neoformans* at 30 °C, 37 °C and 39 °C (Supplemental Figure 16). The ability of the strain lacking the CnTps1 IDD to grow at the higher temperatures indicates that the IDD is not required for growth of *C. neoformans* at temperatures similar to the temperatures observed in human hosts.

*C. neoformans tps1*Δ and *C. neoformans TPS1^IDD^*^Δ^ were unable to grow at 37 °C or 39 °C on YPD plates containing 1 M sorbitol (Supplemental Figure 16). However, wild-type *C. neoformans*, *C. neoformans TPS1^IDD^*^Δ^ and the complementation strain grew in the presence of 1 M sorbitol at 30 °C, 37 °C, and 39 °C (Supplemental Figure 16). These data indicate that *C. neoformans tps1*Δ is sensitive to osmotic stress, but the CnTps1-mediated tolerance of osmotic stress does not require the IDD of CnTps1.

### Specificity of CnTps1 catalytic pocket

To assess the substrate specificity of the CnTps1 catalytic pocket, we tested the catalytic activity of CnTps1 in the presence of UDP-Galactose (UDP-Gal). The hydroxyl groups on the 4^th^ carbon are oriented differently in glucose and galactose. In the CaTps1-UDPG structure, all of the extracyclic hydroxyl groups are easily detected, revealing that the O4 hydroxyl hydrogen forms a hydrogen bond with the peptide backbone of CaTps1 N382 (46). Similarly, the peptide backbone of CnTps1-UDP-G6P residue N517 would interact with the O4 hydroxyl of the glucose moiety of UDPG (Supplemental Figure 17A). Modelling UDP-Galactose into the catalytic pocket reveals the O4 hydroxyl points away from N517, preventing the formation of this hydrogen bond (Supplemental Figure 17B). The loss of this single interaction has a significant effect on the activity of CnTps1 with UDP-Gal as demonstrated by a reduction of only 4% of this substrate and a reduction in catalytic efficiency, compared to CnTps1 with UDPG as a substrate (∼50% use of this substrate) (Figure 6A and Supplemental Table 1), suggesting that a stereochemically proper positioning of the O4, mediated by its interaction with N517, is a critical component of CnTps1 substrate-assisted catalysis.

**Figure 6.**
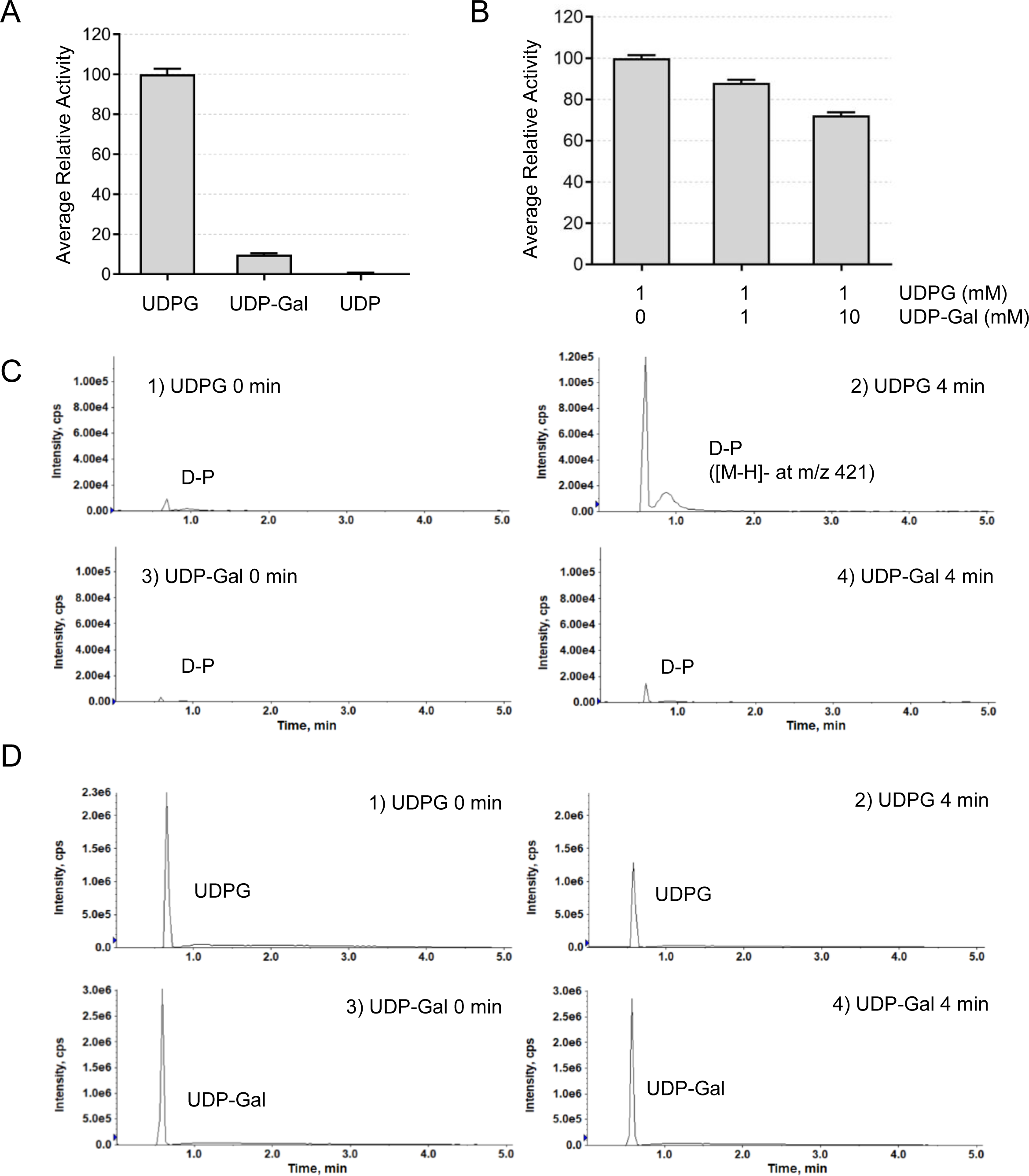
The CnTps1 substrate-binding pocket is highly specific for UDPG. **A)** Relative activity of wild-type CnTps1 utilizing either 1 mM UDPG or 1 mM UDP-Gal or UDP as substrates. Error bars represent the standard error of three independent measurements. **B)** Relative activity of wild-type CnTps1 with 1 mM UDPG and increasing concentrations of UDP-Gal. Error bars represent the standard error of three independent measurements. 1 mM G6P is present in all experiments. **C)** Shown are the extracted ion chromatograms of [M-H]^-^ at *m/z* 421.08 for the formation of a disaccharide-phosphate (D-P). **D)** Shown are the extracted ion chromatograms of [M-H]^-^ at *m/z* 565.05 for UDPG and UDP-Gal at zero and four minutes post catalysis.

To test the effect of UDP-Gal on CnTps1 activity further, we performed a competition assay. The combination of equal molar ratios of UDPG and UDP-Gal to CnTps1 resulted in a 12% reduction in activity, compared to only adding UDPG to the assay (Figure 6B). Interestingly, the addition of 10-fold excess UDP-Gal compared to UDPG resulted in a reduction of CnTps1 activity by approximately 30% (Figure 6B), indicating that UDP-Gal is capable of binding CnTps1 but is a significantly weaker binder and a poor competitive inhibitor. Further, we confirmed that UDP-Gal is a poor substrate of CnTps1 using an orthogonal mass spectrometry-based assay to detect the formation of a disaccharide-phosphate after incubation of CnTps1 with either UDPG or UDP-Gal and the loss of these substrates over 4 minutes (Figure 6C,D). Incubation of CnTps1 with UDPG resulted in the formation of more disaccharide-phosphate (T6P) compared to UDP-Gal (Figure 6C). Consistent with these results, more UDPG was expended in this assay compared to UDP-Gal (Figure 6D). In summary, these results confirm the strong preference of CnTps1 for UDPG.

## Discussion

In order to develop the trehalose biosynthesis pathway as an antifungal drug target, we must improve our understanding of the structure and function of trehalose biosynthesis proteins. Here we report the use of cryo-EM to determine the structures of unliganded CnTps1 and CnTps1 bound to UDP and G6P.

We confirmed that CnTps1 forms a tetramer in solution. Our current hypothesis is that trehalose biosynthesis proteins also form hetero-tetrameric complexes. Interactions amongst Tps proteins (Tps1, Tps2, Tps3 and Tsl1) have been demonstrated in *S. cerevisiae* (63, 64). The function of these complexes may be to sequester the highly cytotoxic trehalose 6-phosphate (T6P) from the cytoplasm, to target trehalose biosynthesis to a proper subcellular localization (65) or to allow for quick and efficient production of trehalose when fungal pathogens are exposed to environmental stress. Future work will include confirmation of the formation of heterocomplexes between CnTps1 and Tps2 both *in vitro* and *in vivo* and determination of the structure of the heterocomplex using cryo-electron microscopy.

We also described herein a conformational change of CnTps1 upon binding of UDP and G6P. Comparison of the unliganded CnTps1 and CnTps1-UDP-G6P structures reveals an inward movement of the N-terminus of CnTps1 when bound to ligand. Most of the movement occurs in α-helix 2 and surrounding residues. The closure of the ligand-bound CnTps1 protomer is consistent with substrate-assisted catalysis, the proposed enzymatic mechanism of Tps1 enzymes (46). Consistent with previous reports of the predicted catalytic activity of fungal Tps1 proteins, we also propose that UDPG binds first, followed by G6P. The proximity of the substrates in the substrate-binding pocket of CnTps1 is demonstrated in Supplemental Figure 9. However, we now can see aspects of the catalytic mechanism occurring in the context of the CnTps1 homo-tetramer.

The more ordered CnTps1-UDP-G6P substrate-binding pocket allows us to observe specific interactions between the ligands and CnTps1 substrate-binding residues. The identification of substrate-binding residues, K420, D514, E522 and R453, provides insights into the mechanism of the Tps1 glycosyltransferase as well as demonstrates the key roles of these residues in substrate specificity (Figure 5A-C). Mutation of any of these residues is sufficient to inhibit CnTps1 catalytic activity (Figure 5D,E). Additionally, our determination of substrate-binding variants which lead to a loss of activity of CnTps1 provides the framework for future investigation of a fully closed substrate-trapped cryo-EM structure of the CnTps1 homo-tetramer. This structure will provide fuller information regarding the CnTps1 catalytic mechanism by allowing visualization of CnTps1 bound to its substrates, UDPG and G6P.

The substrate-binding residues identified in the catalytic pocket of Tps1 are conserved amongst fungal pathogens, including *C. albicans* (Supplemental Figure 1 and Supplemental Figure 10B), supporting the proposition that a broader-spectrum antifungal therapeutic can be developed by targeting the catalytic pocket of Tps1. Additionally, we have been able to demonstrate that the CnTps1 substrate-binding domain is highly specific as underscored by its significantly reduced ability to utilize UDP-Galactose as a substrate, an epimer of UDPG (Figure 6 and Supplemental Figure 17).

Although the key interfaces of the CnTps1 homo-tetramer are not as highly resolved as the substrate-binding pocket, specific residues that may play a role in the formation of the homo-tetramers were identified (Supplemental Figure 12). It remains to be determined whether oligomerization of CnTps1 is required for catalytic activity. If so, in addition to targeting the substrate-binding pocket of CnTps1, the interfaces critical to the formation of the CnTps1 tetramer may also be novel antifungal drug targets.

Finally, we have studied the importance of a 92-residue (residues M209-I300) unstructured region in CnTps1, now called the intrinsically disordered domain (IDD) (Supplemental Figures 1 and 13A). Intriguingly, the IDD is well-conserved amongst Cryptococcal species and closely related fungi in the Basidiomycota division (Supplemental Figure 13B). We have demonstrated, with the generation of a CnTps1 ΔIDD protein, that these 92 residues are not required for the catalytic activity of CnTps1 *in vitro* (Supplemental Figure 13C). These data are consistent with the residues not interacting with CnTps1, neither intra nor intermolecularly, or being proximal to the catalytic region of CnTps1. Indeed, the IDD does not affect the *in vitro* catalytic efficiency of CnTps1 significantly (Supplemental Table 1).

CnTps1 is required for the growth of *C. neoformans* at 37 °C (27). Surprisingly we demonstrated that the CnTps1 IDD is not critical for the survival of *C. neoformans* at higher temperatures and in the presence of osmotic stress, both of which are encountered when *C. neoformans* transitions from the environment to the human host (Supplemental Figure 16). We posit, therefore, that a function of this highly conserved IDD may involve its posttranslational modification, ability to interact with other cellular proteins, targeting CnTps1 to specific subcellular localizations in fungal cells or self-association into a condensate (66, 67). Future work to determine the function of the CnTps1 IDD will aid in the development of the CnTps1 IDD as a novel Basidiomycete-specific antifungal drug target.

CnTps1 has the GT-B fold of the retaining glycosyltransferase family which is characterized largely by a large substrate-binding cleft between two lobes (68). GT-B fold, retaining glycosyltransferases function in a metal-independent manner (68). Several structures of proteins in the GT-B glycosyltransferase family including sucrose synthase from *N. europaea* have been determined (69). While these enzymes have some characteristics that are similar, there are also key differences that make them unique and the context in which they function is also critical. Therefore, to capture the entire wealth of characteristics amongst glycosyltransferases, it is important for new structures to be determined. For example, a comparison of *N. europaea* sucrose synthase and CnTps1 reveals that the key lobes are similar, with an rmsd of 1.4 A (Supplemental Figure 18). However, the sucrose synthase protein has two extra domains (SSN-1, SSN-2) that are not found in CnTps1 whereas CnTps1 contains the IDD, which is not found in sucrose synthase (69). Another surprising difference between these two glycosyltransferases is the lack of the strict conservation of the triplet of conserved catalytic residues amongst members of the GT-B retaining glycosyltransferase family (69). Sucrose synthase and other glycosyltransferases contain a conserved Arg-Lys-Glu triad, in CnTps1 the Glu is replaced by the smaller acidic amino acid residue Asp (69).

In summary, we have determined the first cryo-EM structures of a fungal Tps1 homo-tetramer in its unliganded and ligand-bound conformation. In the future, we shall determine whether a tetramer is potentially required for efficient trehalose biosynthesis and thus, disrupting the oligomerization state of Tps1 might make a viable antifungal drug target (Supplemental Figure 19). We identified key residues that play a role in tetramerization, as well as conserved substrate-binding residues. We have demonstrated that the substrate-binding site of Tps1 is conserved and specific for UDPG. Finally, we identified a 92-residue IDD in CnTps1 that does not contribute to *C. neoformans* Tps1-mediated tolerance of temperature and osmotic stress, both of which are encountered in the human host. However, the conservation of the CnTps1 IDD points strongly towards a necessary function in *Cryptococcus* species as well as Basidiomycetes. In conclusion, our work presented here should facilitate the development of novel antifungal drugs which target multiple aspects of the initial step in the trehalose biosynthesis pathway.

## Materials and Methods

### Bacterial two-hybrid

Protein interactions were analyzed by a bacterial two-hybrid analysis based on a set of vectors described by Daines and Silver (70). The genes for *C. neoformans Tps1* were cloned into the plasmid pSR658, which contains an N-terminal LexA DNA-binding domain. The resulting plasmid was cloned into *E. coli* strain SU101 to test for homo-oligomerization and expression of the protein fusions was induced with 1 mM IPTG. For this purpose, 5 mL of LB culture supplemented with antibiotics were inoculated and grown to 0.5 to 0.6 OD600 at 37 °C. The culture was mixed with permeabilization buffer (100 mM Na_2_HPO_4,_ 20 mM KCl, 2 mM MgSO_4,_ 0.8 mg/mL CTAB, 0.4 mg/mL deoxycholic acid sodium salt, and 5.4 µL/mL of β-mercaptoethanol). Samples were incubated at 30 °C for 30 minutes and subsequently incubated with the prewarmed substrate solution (60 mM Na_2_HPO_4,_ 40 mM NaH_2_PO_4,_ 1 mg/mL ONPG, 2 µL/mL of β-mercaptoethanol). After 25 minutes, the stop solution (1 M Na_2_CO_3_) was added, and the absorbance of the culture supernatant was measured at 420 nM using a SpectraMax M5 plate reader (Molecular Devices).

### CnTps1 Protein expression and purification

The full-length *TPS1* gene from *Cryptococcus neoformans* strain H99 was codon-optimized for expression in *E. coli* (Genscript) and subcloned using ligation-independent cloning into pMCSG7 (71). The construct was transformed into *E. coli* OverExpress C41(DE3) chemically competent cells engineered for high protein expression. Supernatants from lysed cultures, induced with 0.5 mM IPTG, were loaded onto a nickel column (Ni-NTA, Qiagen) and washed in a buffer (50 mM Tris pH 8.0, 300 mM NaCl, 5 mM MgCl_2_ and 5% glycerol) and eluted with increasing amounts of imidazole, which ranged from 40 mM to 400 mM imidazole, in this buffer. The fractions containing 6xHis-CnTps1 were then pooled, concentrated to 5 mL, and run over a S200 size exclusion column (HiLoad 26/600 Superdex 200pg, Cytiva) in pre-cooled buffer containing 20 mM Tris pH 8.0, 300 mM NaCl, 5% glycerol and 2 mM β-mercaptoethanol. Size exclusion fractions containing 6xHis-CnTps1 were pooled and concentrated to 1 mg/mL for downstream experimental applications such as cryo-electron microscopy and activity assays.

### CnTps1 Cryo-EM grid preparation

6xHis-CnTps1 was purified as described above and concentrated in a buffer containing 20 mM Tris pH 8.0, 300 mM NaCl, 5% glycerol and 2 mM β-mercaptoethanol. For grids prepared with unliganded CnTps1, 3 µL of 0.75 mg/mL 6xHis-CnTps1 was applied to glow-discharged carbon Quantifoil grids. After a 15 s incubation, the grids were blotted for 2 s to remove excess protein and rapidly plunged into liquid ethane (−182 °C) using a Leica EM GP2 (Leica Microsystems) operated at 95% humidity and 22 °C. For determination of the structure of the CnTps1-UDP-G6P complex, 0.5 mg/mL 6xHis-CnTps1 was incubated with 10 mM uridine diphosphate (UDP, Sigma) and 10 mM glucose-6-phosphate (G6P, Sigma) for 18 hours at 4 °C. 3 µL of the CnTps1-UDP-G6P mixture was deposited onto glow-discharged UltrAuFoil grids. After a 15 s incubation, the grids were blotted for 2 s to remove excess protein and rapidly plunged into liquid ethane (−182 °C) using a Leica EM GP2 (Leica Microsystems) operated at 95% humidity and 22 °C. Grids were transferred to liquid nitrogen for storage until data collection.

### CnTps1 cryo-EM data collection

After screening for high grid quality, using a Talos Arctica cryo-electron microscope (Thermo Fisher Scientific), a total of 2,520 micrographs from the cryo-EM grids containing unliganded CnTps1 were collected on a Titan Krios cryo-electron microscope (Thermo Fisher Scientific), at 300 kV equipped with a K3 detector (Gatan) at a nominal magnification of 105,000x and defocus values from −2.5 µm to −0.8 µm. The pixel size was 0.65 Å. The total dose was 62 e^−^□Å^−2^.

The cryo-EM grids containing CnTps1-UDP-G6P were also screened on a Talos Arctica cryo-electron microscope (Thermo Fisher Scientific). A total of 3,182 micrographs of CnTps1-UDP-G6P were collected on a Titan Krios cryo-electron microscope (Thermo Fisher Scientific), at 300 kV equipped with a K3 detector (Gatan) at a nominal magnification of 81,000x and defocus values from −2.5 µm to −0.8 µm. The pixel size was 1.08 Å. The total dose was 62 e^−^□Å^−2^.

### Cryo-EM data processing

For unliganded CnTps1, 2,520 dose-fractionated image stacks were aligned using the drift correction routines implemented in RELION3.0 (72) and the contrast transfer function was determined on the motion corrected non-dose-weighted sum of frames using CTFFIND4.1 (73). CnTps1 particles were boxed out using template-free Auto-picking in RELION3.0. Two consecutive rounds of 2D classification were preformed to obtain a clean set of particles. The initial model was generated in RELION3.0, and 3 rounds of 3D classification were performed with resulting reconstructions showing non-uniform angular distributions. To improve the alignment, 907 micrographs were selected with the CTF max fitting resolution cutoff as 4 Å. A total of 456,020 4x binned clean particles were used for further processing. A new *ab initio* 3D reference was generated using 16,000 particles followed by 3D classification, and particles were classified into 5 classes. 207,081 particle images corresponding to best class were kept and subjected to 3D refinement without symmetry resulting in a 5.3 Å resolution map. Particles were then re-centered and re-extracted without binning and further refined using a shape mask to ∼3.6 Å resolution. After CTF refinement and Bayesian polishing in RELION3.0, the polished particles were imported into cryoSPARC (74). Homogeneous refinement was performed without imposing symmetry resulting in resolution of 3.6 Å. After imposing D2 symmetry, the final global resolution of the reconstruction improved to 3.3 Å. A summary of the data processing steps is shown in Supplemental Figure 3.

For determination of the structure of CnTps1 bound to UDP-G6P, dose-fractionated movies were aligned with MotionCor2 (75) and CTF estimation was performed using CTFFIND4.1. Aligned micrographs were sorted and selected according to max fitting resolution with a cutoff of 4 Å with cryoSPARC Curate Exposures yielding 2,617 images. CnTps1 particles were boxed out with template-free Blob picker. Approximately, 2.8 million particles were extracted and subjected to 2D classification using a circular mask (180 Å radius) to focus the alignment on one protomer. A total of 2,079,376 clean particles were selected and subjected to *ab initio* Reconstruction using 4 classes. A conformationally homogeneous class showed well resolved features and accounted for ∼850k particles which were kept and subjected to 3D refinement (without imposing symmetry) resulting in a 3.1 Å resolution map. To improve the density of the flexible loops around the substrate binding pocket, 3D variability analysis was conducted using cryoSPARC, and the particles were classified into 8 clusters. Further non-uniform refinement using only particles assigned to cluster 1 resulted in the best map (3.5 Å resolution) that showed well-resolved features on the flexible loop region surrounding the catalytic pocket. A summary of the data processing steps is shown in Supplemental Figure 6.

### Cryo-EM model building

Model building of the unliganded CnTps1 structure was initiated by using the CaTps2 N-terminal domain crystal structure (PDB ID 5XDF) as the starting model. α-helix 2 had to be rebuilt. CnTps1 residues were mutated to match the correct sequence using COOT (76). Coordinates were then fitted manually in Coot (76) followed by iterative refinement using Phenix (77) real space refinement to improve the quality of the models. Following completion of the model of a single protomer, multiple copies of the models were generated and docked into the map, using UCSF Chimera 1.14, to form the homo-tetramer (78). The CnTps1-UDP-G6P model was generated using the same workflow. The exception is that the initial model of the complex was the unliganded CnTps1 model.

### Tps1 Activity Assay

The catalytic activity of Tps1 was measured utilizing a continuous enzyme coupled assay as previously reported (79). Briefly, 6xHis-CnTps1 protein was concentrated in a buffer containing 20 mM Tris pH 8.0, 300 mM NaCl, 5% glycerol and 2 mM β-mercaptoethanol. The assay was carried out in a buffer containing 50 mM HEPES, pH 7.8, 100 mM KCl, 5 mM MgCl_2_ and 2 mM DTT. A final concentration of 3 μM, 1 mM and 1 mM were utilized for Tps1, uridine diphosphate glucose and glucose-6-phosphate, respectively. When reported, UDP-Galactose (UDP-Gal, Sigma) was used in the assay. Activity assays were performed in clear, flat-bottomed 96-well plates and the decrease in absorbance at 340 nm was recorded using a SpectraMax M5 plate reader (Molecular Devices). The decrease in absorbance at 340 nm was analyzed for the 200 s of the reaction, which corresponds to the initial rate of the reaction.

### Circular Dichroism spectroscopy

Far-UV CD spectra of CnTps1 ΔIDD were recorded on an AVIV 435 CD Spectrophotometer in a 1□mm sample cell. Measurements were taken from 200 to 260□nm with a wavelength step of 1.0□nm and a 1□s averaging time. Each spectrum is the average of 3 scans. CnTps1 ΔIDD was buffer exchanged into CD buffer (20□mM NaH_2_PO_4_ (pH 7.5), 300□mM NaF and 5% glycerol) and concentrated to a final concentration of 0.5 mg/mL.

### Generation of CnTps1 strains

Strains and primers used in this study are listed in Supplemental Table 2. The following strains were constructed in this study: *tps1*Δ, *TPS1^IDD^*^Δ^, and *TPS1^IDD^*^Δ^*::TPS1.* To construct the *tps1*Δ deletion mutant strain, three PCRs were prepared: the 5’ flanking region (AD131/AD132), the nourseothricin (NAT) drug selection cassette (AD133/AD134) amplified from plasmid, pIA3 (80), and the 3’ flanking region (AD135/AD136). These PCR fragments were gel extracted, purified using the Qiaquick gel extraction kit (Qiagen), and cloned into linearized pUC19 by In-Fusion cloning (Takara Bio). To construct the *TPS1^IDD^*^Δ^ strain, four PCRs were prepared: the 5’ flank region (AD3710/AD3711), the neomycin (NEO) drug selection cassette amplified from plasmid, pJAF1 (81) (AD3712/AD3713), 5’ flank region closest to *TPS1* and the *TPS1* gene up to the IDD (AD3714/AD3715), and part of the *TPS1* gene downstream of the IDD (AD3716/AD3717). Primers AD3715 and AD3716 have the Ca IDD nucleotide sequence codon optimized for *Cryptococcus* for replacement of the CnTPS1 IDD. These fragments were cloned into linearized pUC19 using the Hi-Fi Assembly Cloning Kit (New England Biolabs). The *tps1*Δ mutant was confirmed by PCR and the TPS1*^IDD^*^Δ^ strain was confirmed by PCRs and sequencing to confirm the replacement of the CnTps1IDD with the CaTps1IDD. Lastly, a wild-type copy of *TPS1* was introduced into the *TPS1^IDD^*^Δ^ strain. Three PCRs were performed to amplify: the 5’ flanking region (AD3748/AD3749), the nourseothricin drug selection cassette (NAT) (AD3750/AD3751), and additional 5’ flank region and part of the *TPS1* gene (AD3572/AD3573). These fragments were cloned into linearized pUC19 using the Hi-Fi Assembly Cloning Kit (New England Biolabs). The plasmids carrying the desired inserts were transformed into strains by biolistic transformation as previously reported (82). Strains were confirmed by PCR using primers listed in Supplemental Table 2 and sequencing.

### LC/MS/MS

Samples for LC/MS/MS were prepared by incubating by 2 µM CnTps1 with 1 mM G6P and either 1 mM UDPG or 1 mM UDP-Gal for 4 minutes in a reaction buffer consisting of 20 mM Tris pH 8.0, 300 mM NaCl, 5% glycerol and 2 mM β-mercaptoethanol. The reactions were stopped by the addition of methanol. Samples were centrifuged at 10,000 rpm at room temperature for 2 minutes. Subsequently, the supernatant was collected and submitted for analysis.

Reverse-phase liquid chromatography-electrospray ionization/tandem mass spectrometry (LC-ESI/MS/MS) was performed using a Shimadzu LC system (comprising a solvent degasser, two LC-10A pumps and a SCL-10A system controller) coupled to a high-resolution TripleTOF5600 mass spectrometer (AB Sciex, Framingham, MA). LC was operated at a flow rate of 200 μl/min with a linear gradient as follows: 100% of mobile phase A was held isocratically for 2 min and then linearly increased to 100% mobile phase B over 5 min and held at 100% B for 2 min. Mobile phase A was a mixture of water/acetonitrile (98/2, v/v) containing 0.1% acetic acid. Mobile phase B was a mixture of water/acetonitrile (10/90, v/v) containing 0.1% acetic acid. A Zorbax SB-C8 reversed-phase column (5 μm, 2.1 x 50 mm) was obtained from Agilent (Palo Alto, CA). The LC eluent was introduced into the ESI source of the mass spectrometer. Instrument settings for negative ion ESI/MS and MS/MS analysis of lipid species were as follows: Ion spray voltage (IS) = −4500 V; Curtain gas (CUR) = 20 psi; Ion source gas 1 (GS1) = 20 psi; De-clustering potential (DP) = −55 V; Focusing potential (FP) = −150 V. Data acquisition and analysis were performed using the Analyst TF1.5 software (AB Sciex, Framingham, MA).

## Data, Materials, and Software Availability

Cryo-EM density maps of unliganded CnTps1 and CnTps1-UDP-G6P have been deposited in the Electron Microscopy Databank (EMDB) (https://www.ebi.ac.uk/emdb) with accession codes EMD-29338 and EMD-29172, respectively (83, 84). Atomic models of unliganded CnTps1 and CnTps1-UDP-G6P have been deposited in the RCSB Protein Data Bank (PDB) (https://www.rcsb.org) with accession codes PDB ID 8FO1 and PDB ID 8FHW, respectively (85, 86). The enzymatic data in this study have been deposited in the Standards for Reporting Enzymology Data (STRENDA) database with the SRN HSMHV7 and VZVQFC. All software used in this study is publicly available. The paper does not report any original code. All other data are included in the manuscript and/or supplemental data.

## Acknowledgements

This work was funded by grant 1P01AI104533 from the U.S. National Institutes of Health to R.G.B. and the NIH Tri-Institutional Molecular Mycology and Pathogenesis Training Grant T32AI052080 to E.W. Cryo-EM grid preparation and screening were carried out at the Genome Integrity and Structural Biology Laboratory at the National Institute of Environmental and Health Sciences in Research Triangle Park, NC. Cryo-EM data collection was performed at the Duke University Shared Materials Instrumentation Facility (SMIF), a member of the North Carolina Research Triangle Nanotechnology Network (RTNN), which is supported by the National Science Foundation as part of the National Nanotechnology Coordinated Infrastructure (NNCI) (Grant ECCS-2025064). We thank Dr. Mark Walters, Allen Hsu, and Megan Kopp for cryo-EM data collection support. We thank Dr. Brady Travis for cryo-EM data processing support. This study utilized the computational resources offered by Duke Research Computing. The open access option is selected.

## Author Contributions

E.J.W. and R.G.B. designed the experiments and analyzed the biochemical data. E.J.W., Y.Z., A.B., and R.G.B. analyzed the structural data. E.J.W. generated purification constructs, purified proteins, determined their structures and performed biochemical characterizations. J.L.T., D.L.T., and J.R.P. designed and performed cellular experiments. Y.G. performed mass spectrometry experiments. M.P., A.H., Y.Z., M.J.B. and A.B. provided cryo-EM consulting and experimental input. E.J.W., Y.Z. and R.G.B. wrote the manuscript with significant input from A.B. All authors have read and approved the manuscript.

---

**Data Table 1.**
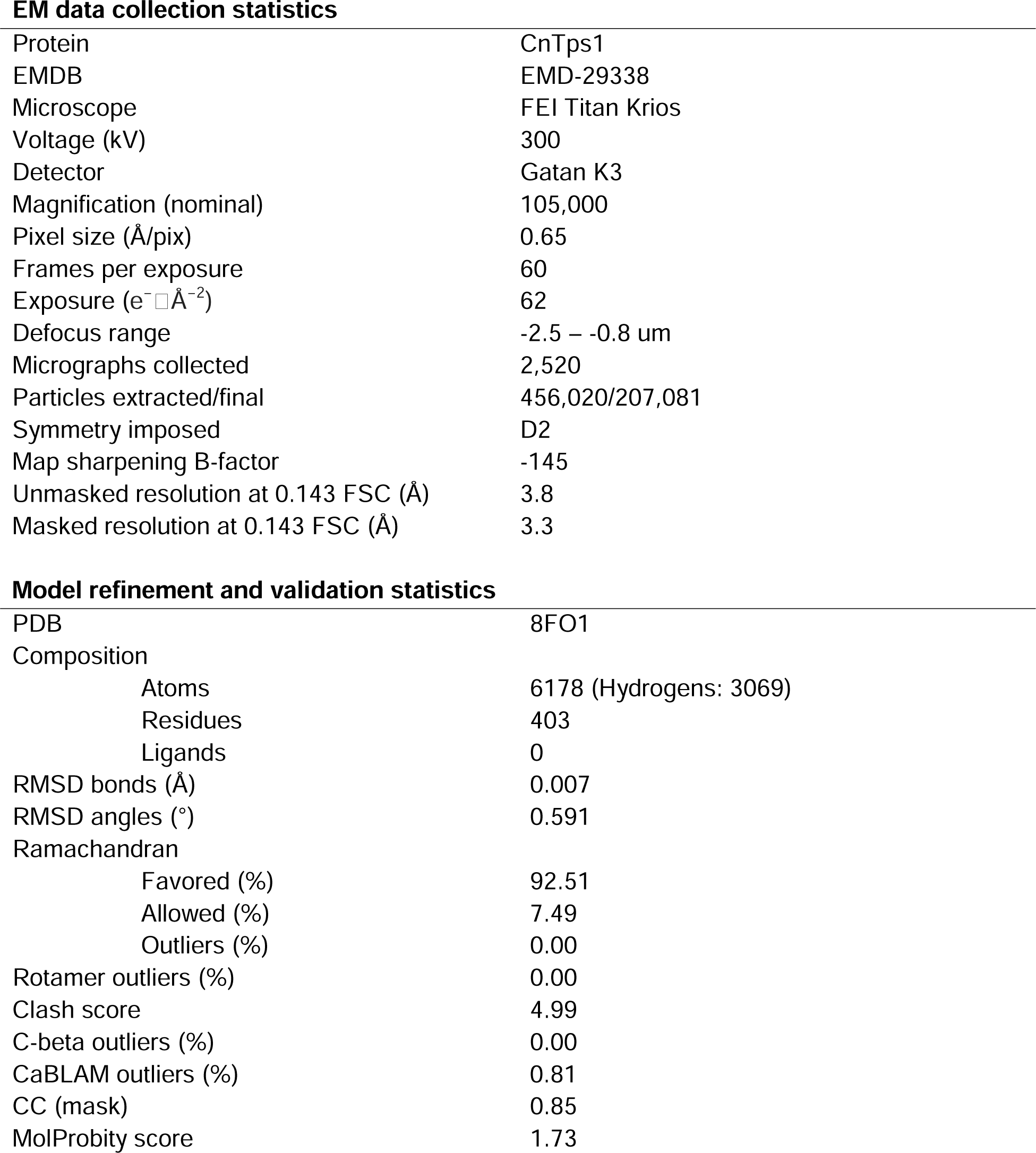
Cryo-EM data collection and refinement statistics for unliganded CnTps1. Data collection and refinement statistics regarding the cryo-EM structure of unliganded CnTps1. FSC, Fourier shell correlation.

**EM data collection statistics**
|  |  |
| --- | --- |
| Protein | CnTps1 |
| EMDB | EMD-29338 |
| Microscope | FEI Titan Krios |
| Voltage (kV) | 300 |
| Detector | Gatan K3 |
| Magnification (nominal) | 105,000 |
| Pixel size (Å/pix) | 0.65 |
| Frames per exposure | 60 |
| Exposure (e<sup>-</sup> Å<sup>-2</sup>) | 62 |
| Defocus range | -2.5 – -0.8 um |
| Micrographs collected | 2,520 |
| Particles extracted/final | 456,020/207,081 |
| Symmetry imposed | D2 |
| Map sharpening B-factor | -145 |
| Unmasked resolution at 0.143 FSC (Å) | 3.8 |
| Masked resolution at 0.143 FSC (Å) | 3.3 |
| **Model refinement and validation statistics** |  |
| PDB | 8FO1 |
| Composition |  |
| Atoms | 6178 (Hydrogens: 3069) |
| Residues | 403 |
| Ligands | 0 |
| RMSD bonds (Å) | 0.007 |
| RMSD angles (°) | 0.591 |
| Ramachandran |  |
| Favored (%) | 92.51 |
| Allowed (%) | 7.49 |
| Outliers (%) | 0.00 |
| Rotamer outliers (%) | 0.00 |
| Clash score | 4.99 |
| C-beta outliers (%) | 0.00 |
| CaBLAM outliers (%) | 0.81 |
| CC (mask) | 0.85 |
| MolProbity score | 1.73 |

**Data Table 2.**
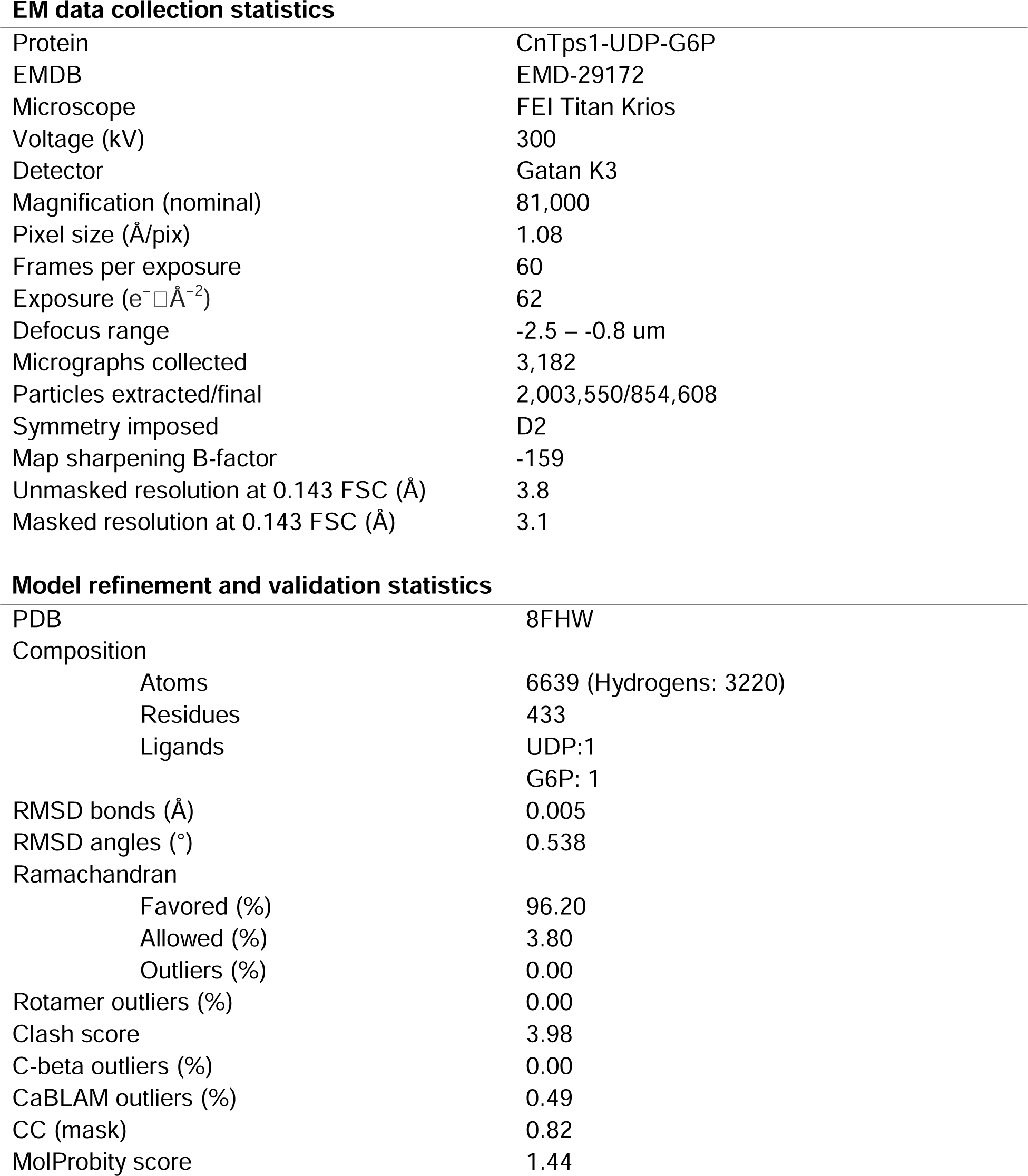
Cryo-EM data collection and refinement statistics for CnTps1-UDP-G6P. Data collection and refinement statistics regarding the cryo-EM structure of CnTps1 in complex with UDP and G6P. FSC, Fourier shell correlation.

