## Supplemental Data for "Structures of trehalose-6-phosphate synthase, Tps1, from the fungal pathogen *Cryptococcus neoformans*: a target for novel antifungals"

**This PDF file includes:**

Figures S1 to S19

Tables S1 to S2

SI References

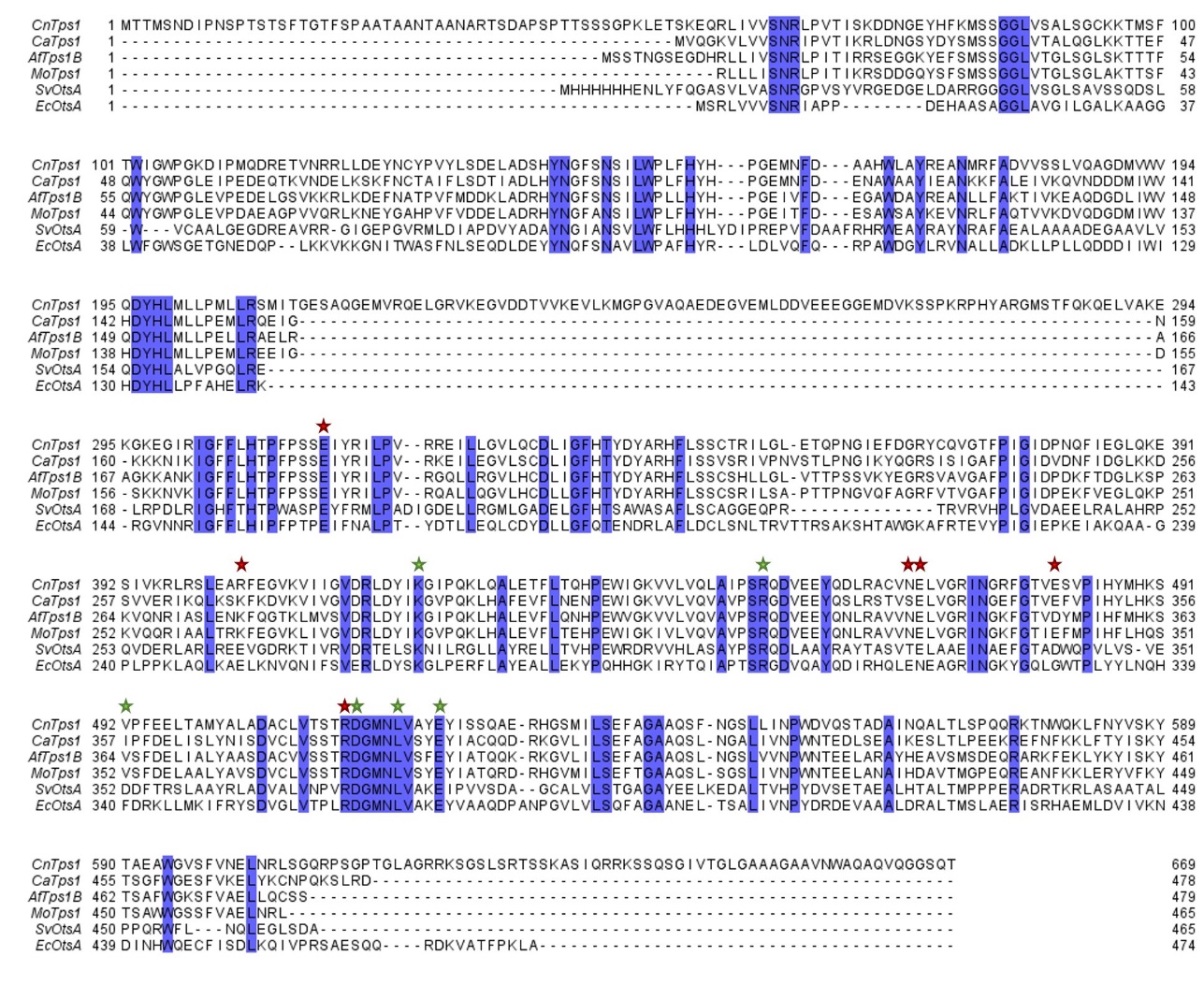

**Figure S1. Sequence-based alignment of Tps1 of *C. neoformans* with other selected species.** Tps1 sequences of *Cryptococcus neoformans*, *Candida albicans*, *Aspergillus fumigatus*, *Magnaporthe oryzae*, *Streptomyces venezuelae*, and *Escherichia coli*. Key substrate-binding residues are highlighted with green stars. Key residues in interfaces between protomers are highlighted with red stars. Unstructured regions specific to *C. neoformans* Tps1 can be visualized in the N-terminus, C-terminus and between residues 209-300. The insertions found in *C. neoformans* Tps1 are not present in the Tps1 proteins from *Candida albicans*, *Aspergillus fumigatus*, *Magnaporthe oryzae*, *Streptomyces venezuelae*, and *Escherichia coli*.

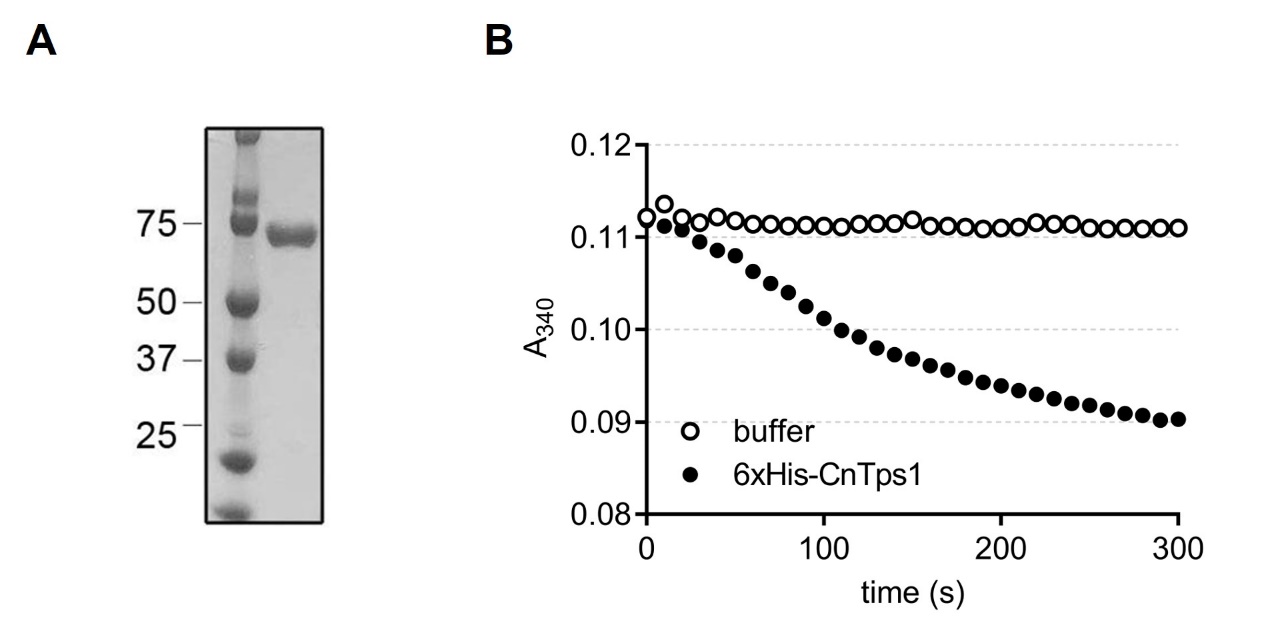

**Figure S2.** **Purification and characterization of 6xHis-CnTps1. A)** Coomassie-stained SDS-PAGE gel demonstrating the purification of 6xHis-CnTps1. 6xHis-CnTps1 was purified via nickel affinity chromatography and followed by size exclusion chromatography. **B)** Spectrophotometric-coupled enzyme assay measuring the activity of 6xHis-CnTps1 as indicated by the decrease in absorbance at 340 nm. CnTps1 enzymatic activity is measured by coupling UDP production and pyruvate kinase activity to the conversion of phosphoenolpyruvate (PEP) to pyruvate and then lactate dehydrogenase (LDH) to convert pyruvate to lactate, which oxidizes NADH to NAD^+^ resulting in a decrease of absorbance at 340 nm.

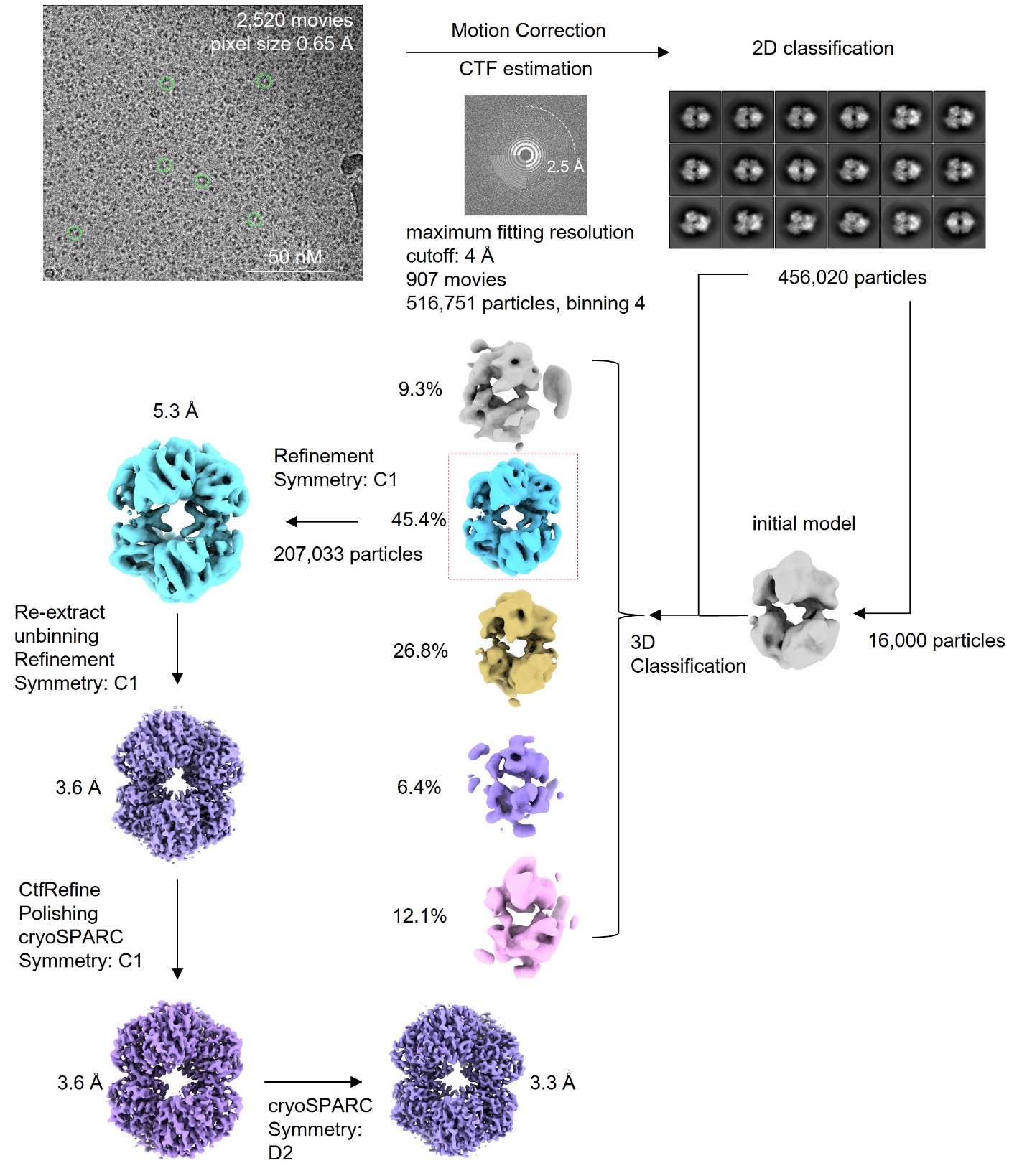

**Figure S3. Cryo-EM data processing workflow of the unliganded CnTps1 dataset.** Single particle analysis workflow used to obtain the high-resolution reconstruction of unliganded CnTps1.

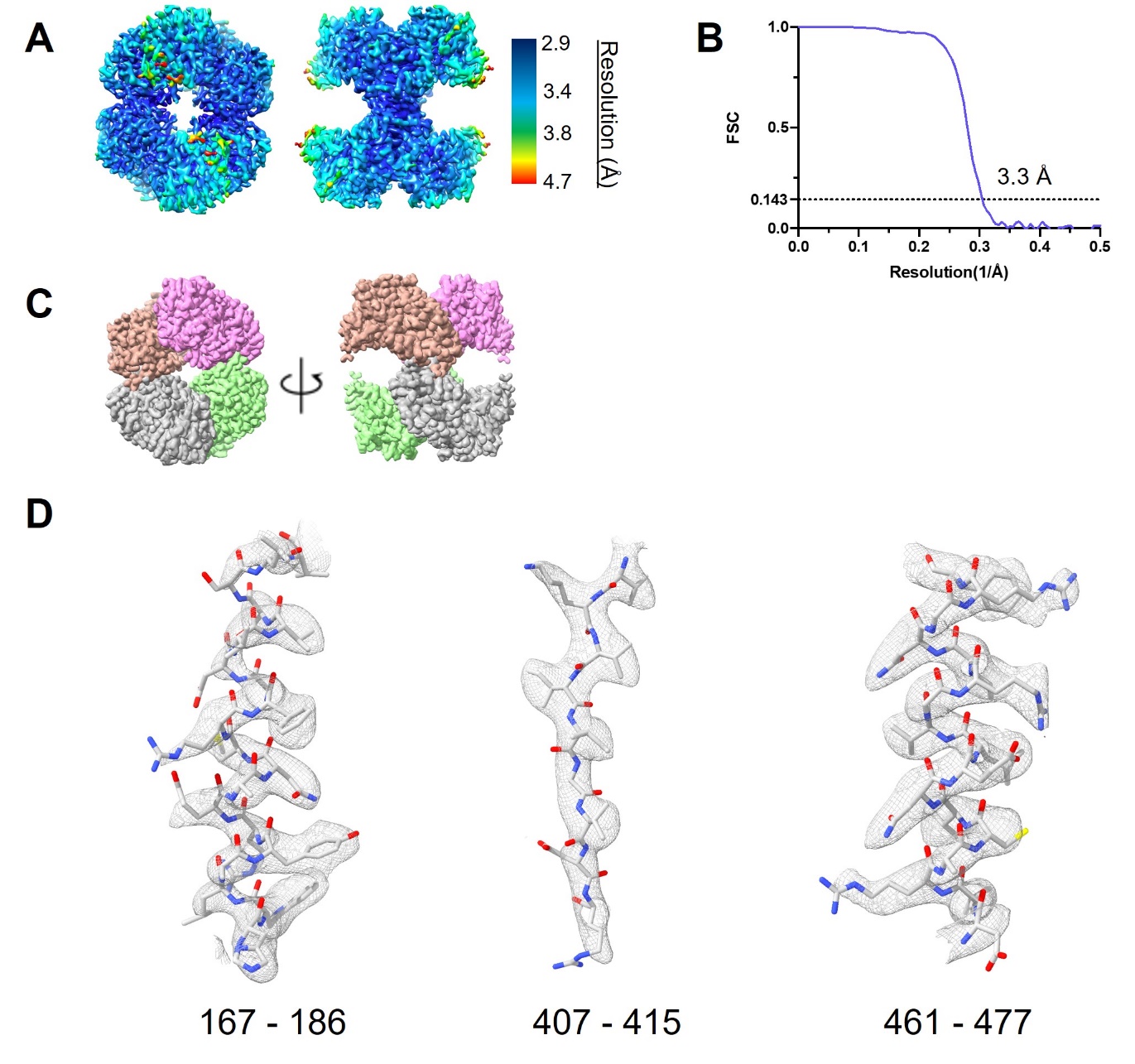

**Figure S4. Unliganded CnTps1 cryo-EM map quality. A)** Final cryo-EM density map colored according to local resolution, shown in Å. **B)** Fourier Shell Correlation (FSC) plot between half-maps showing an estimated map resolution of 3.3 Å (0.143-cutoff criteria). **C)** Space-filling model of unliganded CnTps1 homo-tetramer colored according to protomers and rotated 90° around the vertical axis. **D)** Selected regions of the structure showing the atomic model fit into the cryo-EM map (grey mesh) comprising residues: α-helix 167-186, β-strand 407-415, and α-helix 461-477.

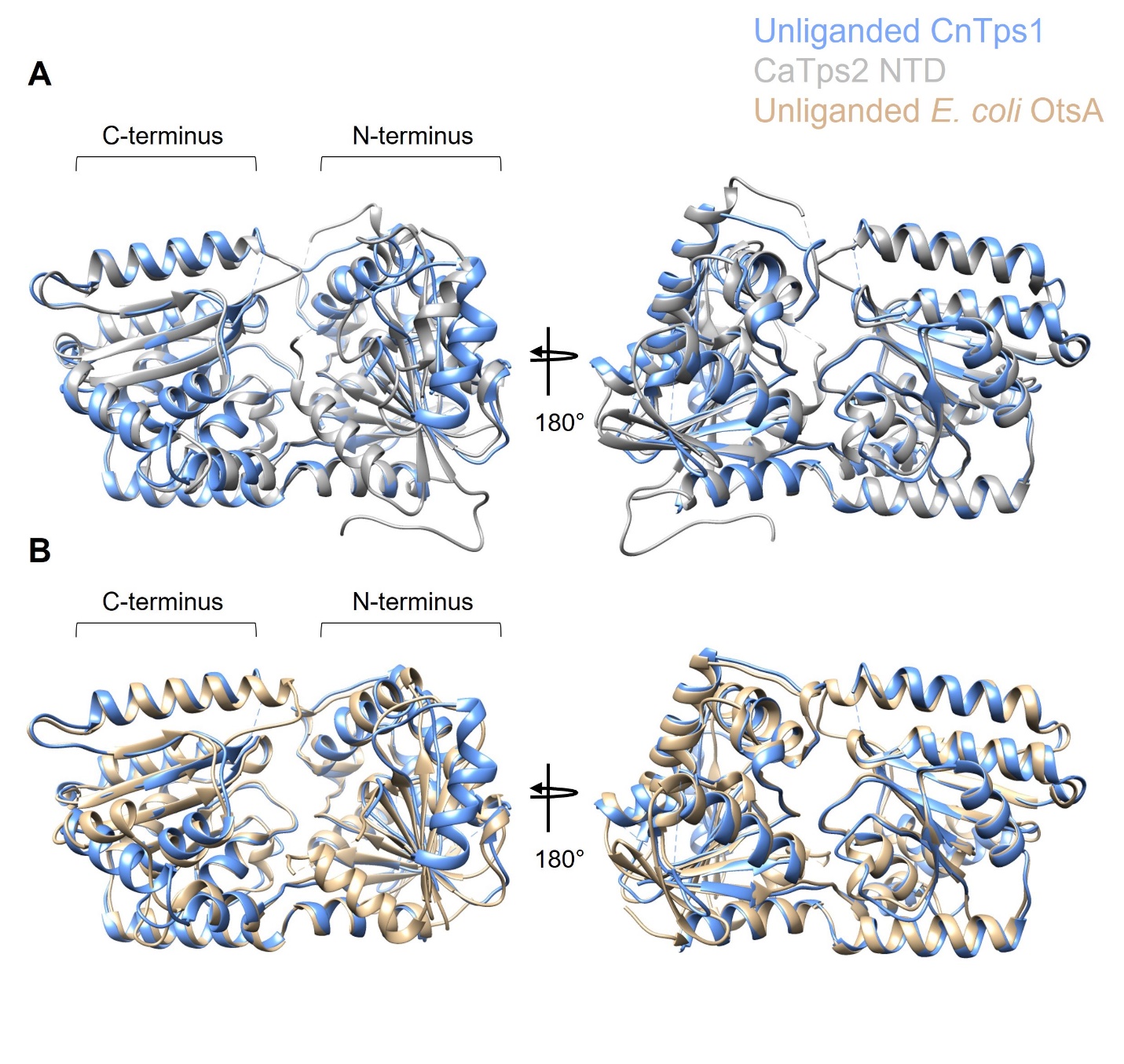

**Figure S5. Comparison of unliganded CnTps1 structures.** **A)** An overlay of an unliganded CnTps1 protomer (in light blue) from the cryo-EM structure with the Tps1-like N-terminal domain of *C. albicans* Tps2 (grey; PDB ID 5DXF) (1). **B)** An overlay of an unliganded CnTps1 protomer (in light blue) and unliganded *E. coli* OtsA (light brown; PDB ID 6JAK) (2). All structures are shown from the “front” and rotated 180°. The N and C-terminal lobes are labelled. Dashed lines represent residues that could not be built into the model.

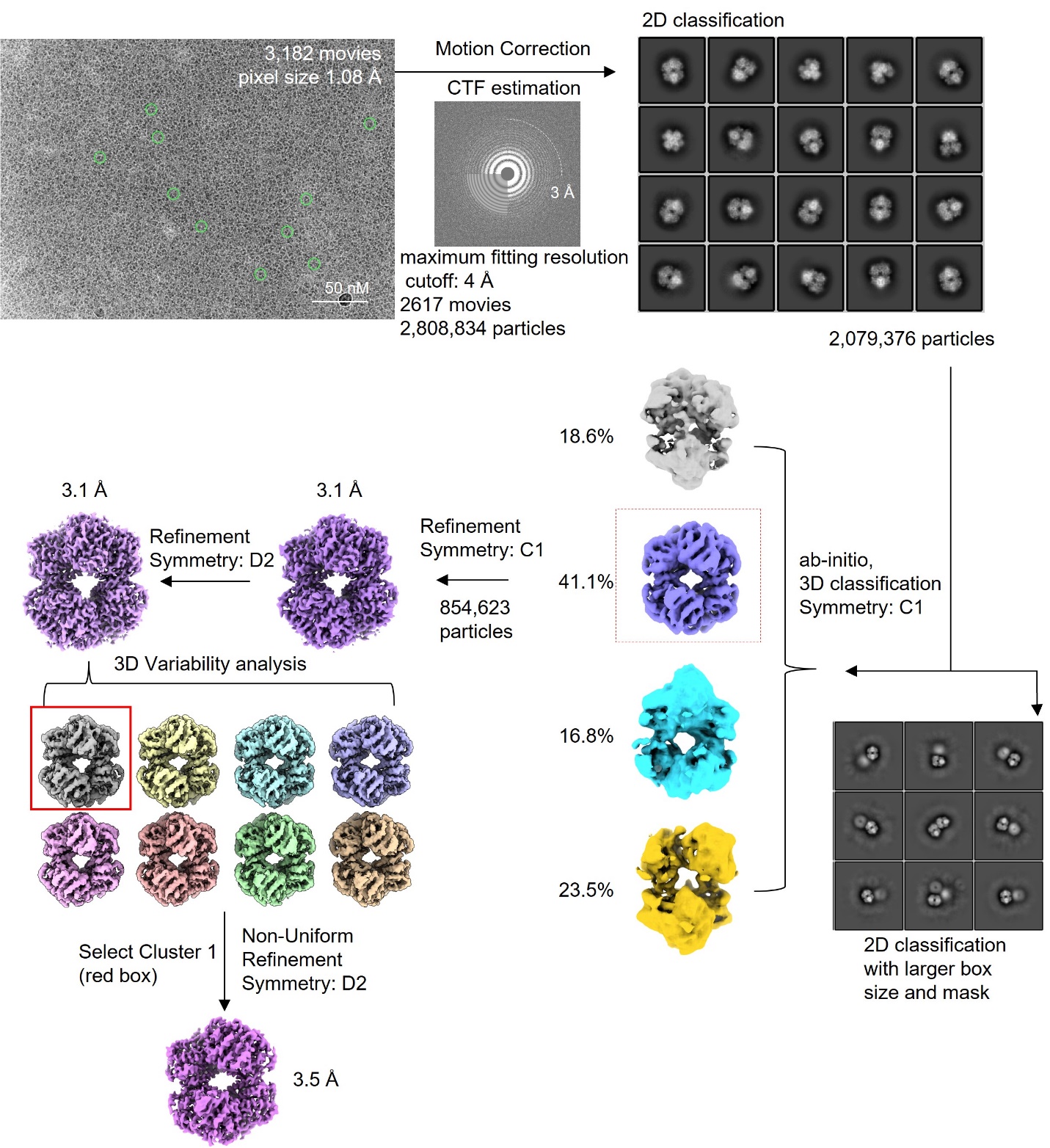

**Figure S6. Cryo-EM data processing workflow of the CnTps1-UDP-G6P dataset.** Single particle analysis workflow used to obtain the high-resolution reconstruction of CnTps1-UDP-G6P.

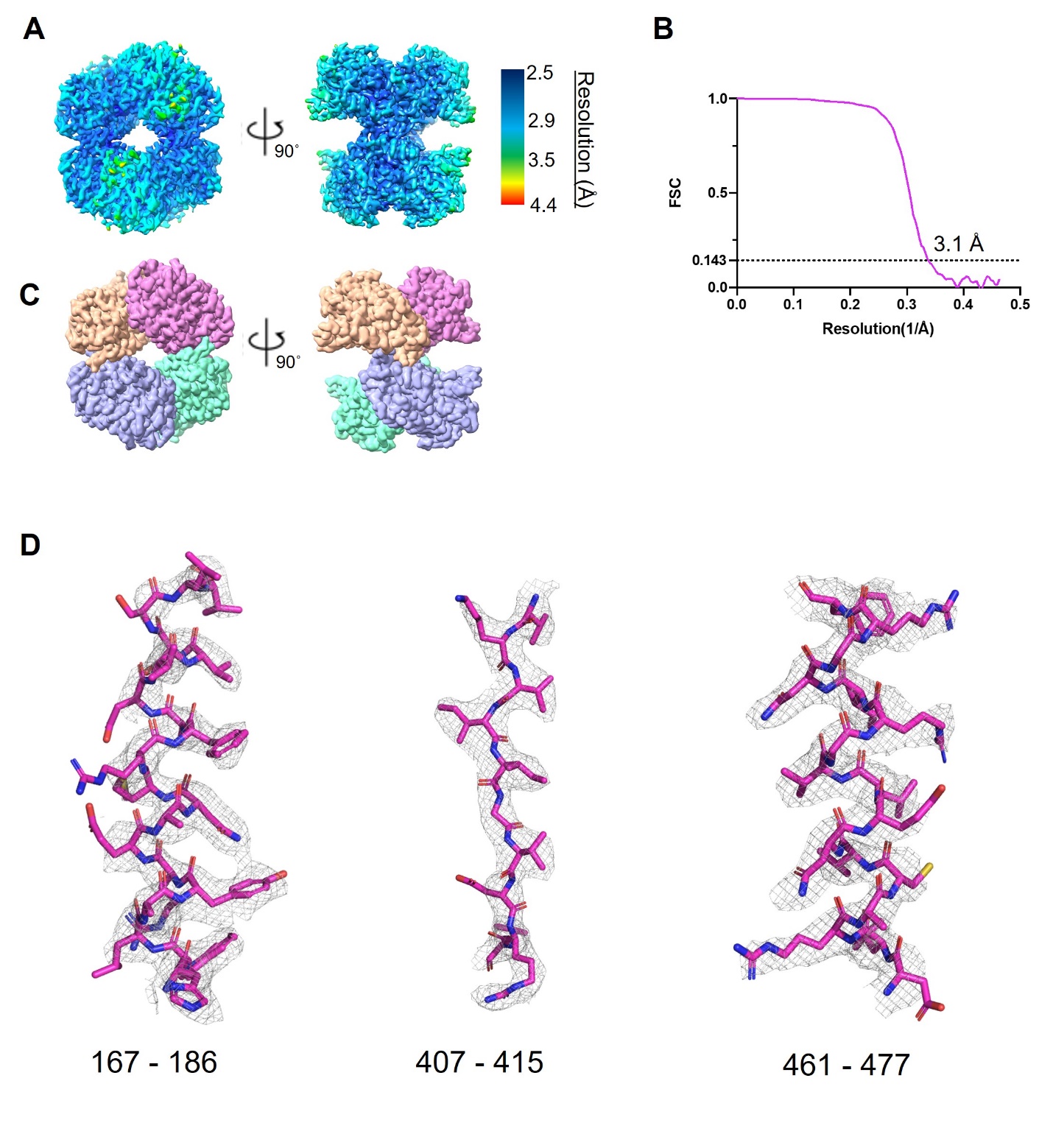

**Figure S7. CnTps1-UDP-G6P map quality.**

**A)** The 3.1 Å consensus cryo-EM density map colored according to local resolution. **B)** Fourier Shell Correlation (FSC) plot between half-maps showing an estimated map resolution of 3.1 Å (0.143-cutoff criteria). **C)** Space-filling model of CnTps1-UDP-G6P homo-tetramer colored according to protomers and rotated 90° around the vertical axis. **D)** Selected regions of the structure showing the atomic model fit into the cryo-EM map (grey mesh) comprising residues: α-helix 167-186, β-strand 407-415, and α-helix 461-477.

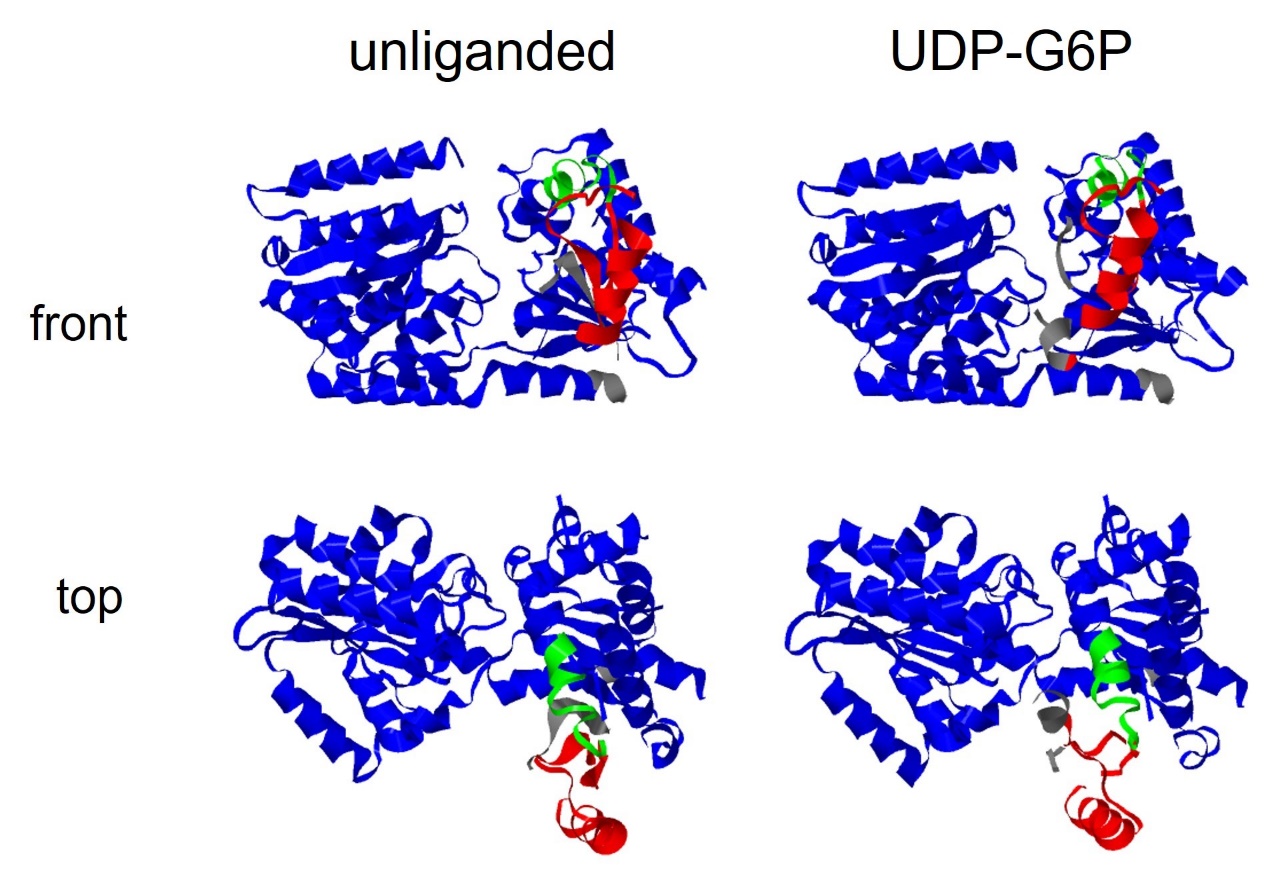

**Figure S8. DynDom analysis revealing the movement of α-helix 2.**

DynDom (3, 4) analysis shows the movement of CnTps1 α-helix 2 (residues 112–125) centered in the flexible region highlighted in red (residues 90-133). The hinge regions (residues 133–145) are displayed in green. Structures of the CnTps1 protomers, both unliganded and UDP-G6P-bound, are shown in ribbon diagrams from top and front views.

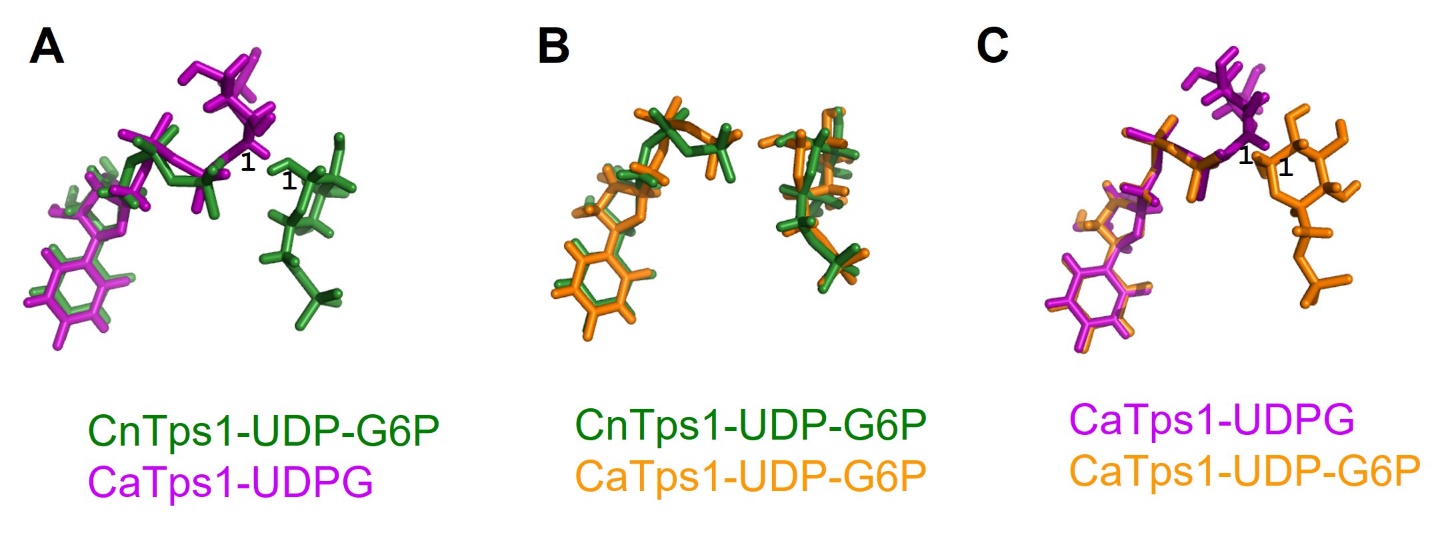

**Figure S9. Comparison of ligands in CnTps1 and CaTps1 structures.**

**A)** An overlay of the ligands from the CnTps1-UDP-G6P cryo-EM structure (green) and CaTps1-UDPG crystal structure (magenta; PDB ID 5HUT). Position 1 on the glucose moieties of UDPG and G6P are labelled. **B)** An overlay of the UDP and G6P molecules from the CnTps1-UDP-G6P cryo-EM structure (green) and CaTps1-UDP-G6P crystal structure (orange; PDB ID 5HUU). **C)** An overlay of ligands from the CaTps1-UDPG (magenta; PDB ID 5HUT) and CaTps1-UDP-G6P (orange; PDB ID 5HUU) crystal structures. Position 1 on the glucose moieties of UDPG and G6P are labelled.

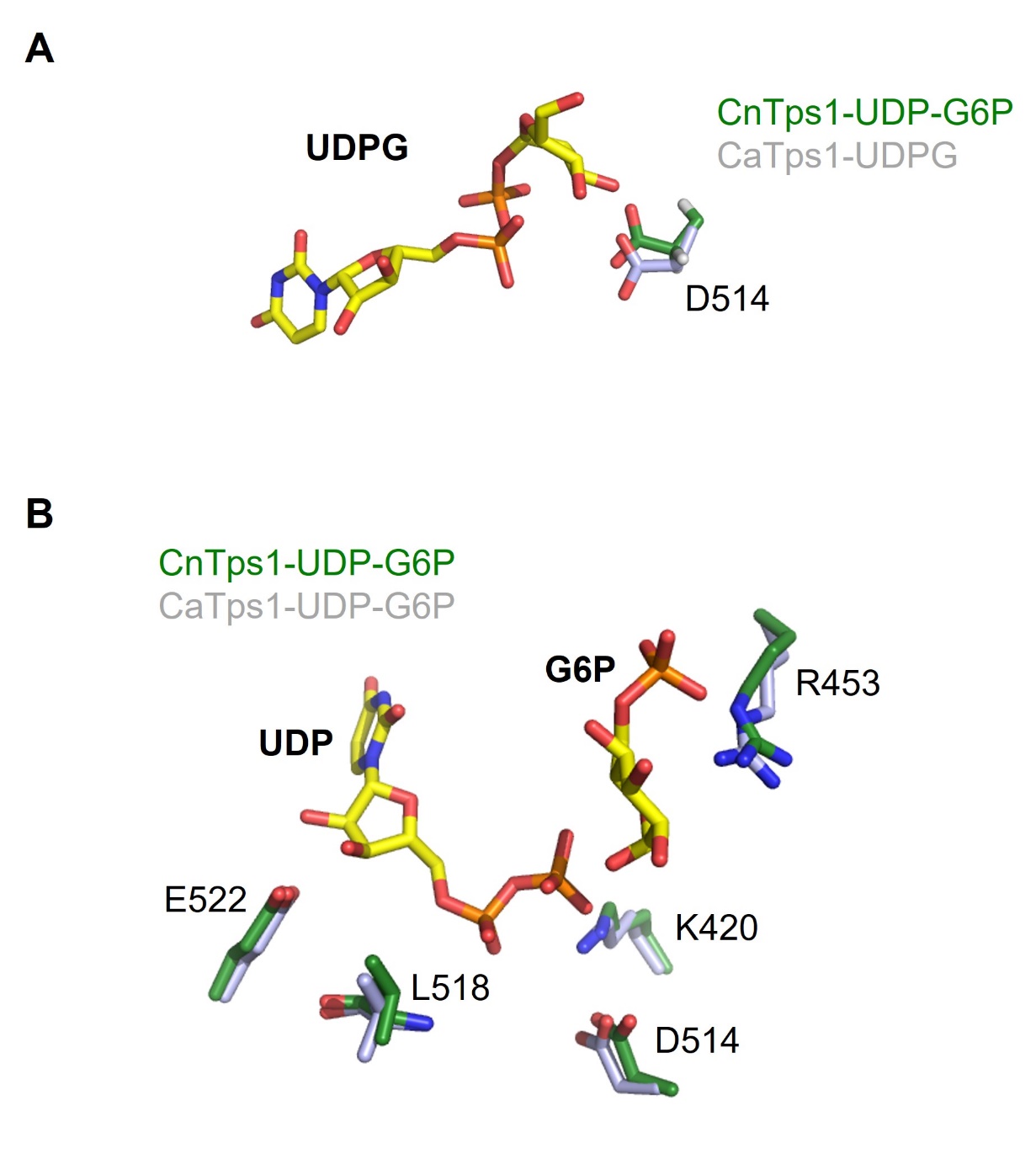

**Figure S10. The interactions of conserved CnTps1 substrate-binding residues with substrates. A)** An overlay of CnTps1-UDP-G6P protomer (green) with CaTps1-UDPG (grey; PDB ID 5HUT). CnTps1 substrate-binding residue D514 is shown as green sticks and it aligns with CaTps1-UDPG residue, D379, shown as grey sticks. The UDPG molecule from CaTps1 is shown as atom-colored sticks. **B)** Overlay of CaTps1-UDP-G6P (PDB ID 5HUU) (shown in grey) substrate-binding residues with CnTps1-UDP-G6P substrate-binding residues (shown in green). The residue numbers for CnTps1-UDP-G6P are labelled. The UDP and G6P molecules from CnTps1-UDP-G6P are shown as atom-colored sticks.

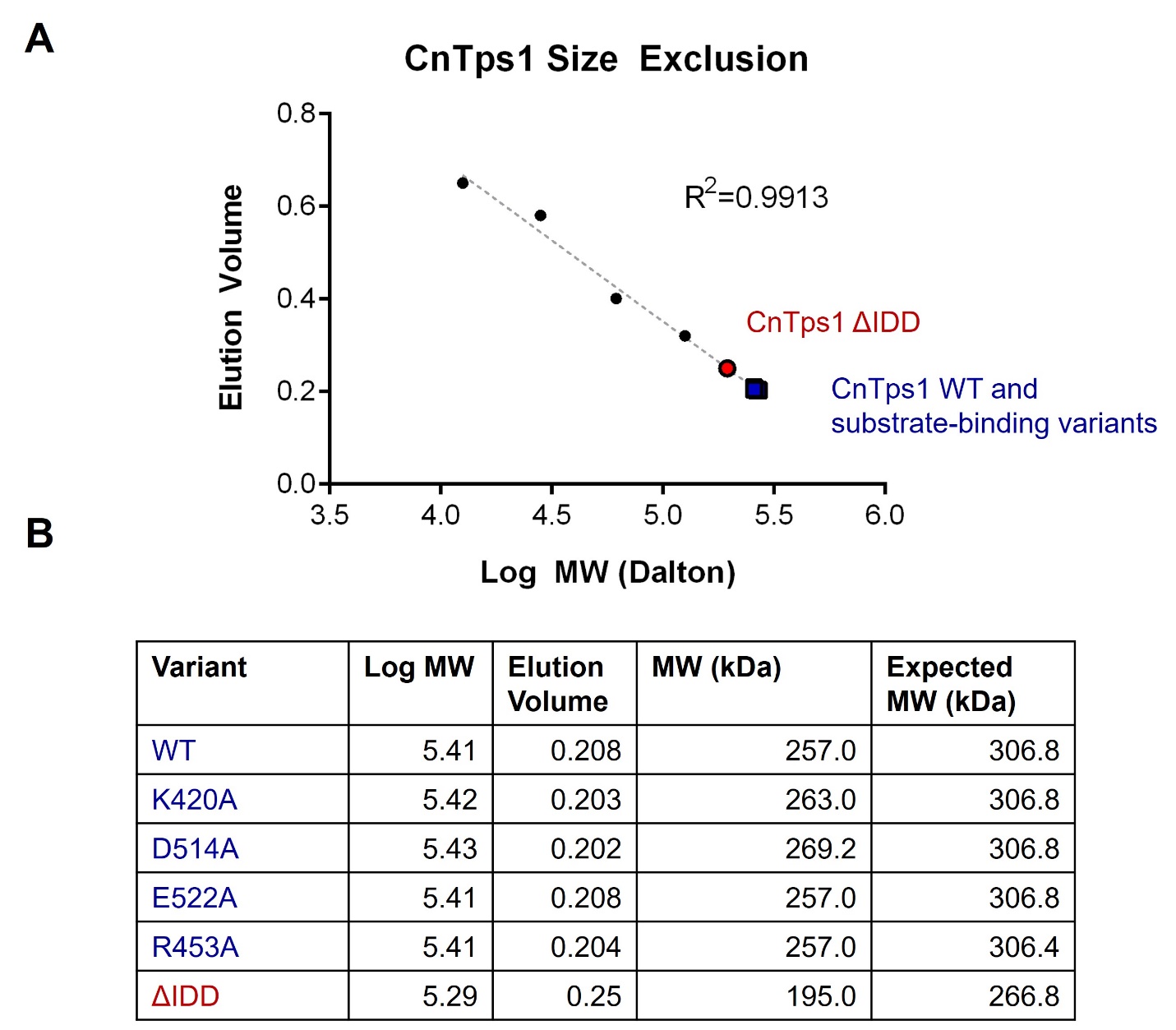

**Figure S11.** **Determination of the apparent molecular mass of wild-type 6xHis-CnTps1 and variants by size exclusion chromatography. A)** Standard curve generated from a linear fit of the log molecular weight of the protein standards (small filled black circles) versus their elution volume, K_av_, using an S200 size exclusion column (HiLoad 26/600 Superdex 200pg, Cytiva). The four standards are cytochrome C (12.5 kDa), carbonic anhydrase (29 kDa), albumin (66.5 kDa) and alcohol dehydrogenase (150 kDa). Log molecular weights of wild-type CnTps1 and variants CnTps1 K420A, D514A, E522A and R453A were plotted versus their normalized elution volume and shown as blue squares. Similarly, the log molecular weight of the CnTps1 ΔIDD was plotted and represented as a red circle. **B)** Table detailing the CnTps1 variants and their respective log molecular weights, elution volumes, molecular weights and expected molecular weights based on sequences.

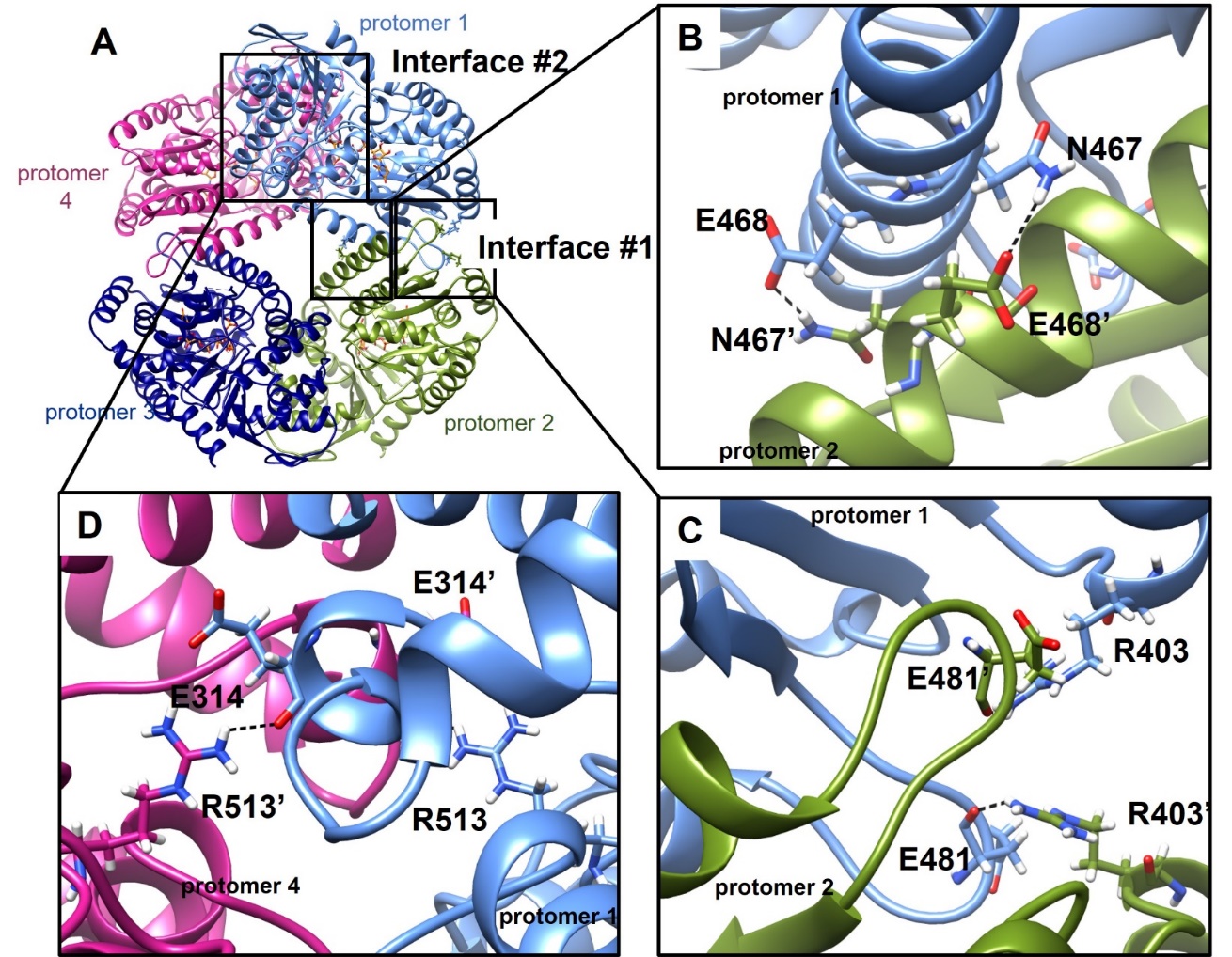

**Figure S12. Interactions that stabilize the CnTps1 homo-tetramer interface. A)** The structure of the CnTps1-UDP-G6P homo-tetramer shown as a ribbon diagram. The four protomers of the CnTps1 homo-tetramer are light blue, green, navy and magenta. The two tetrameric interfaces, labelled Interface #1 and Interface #2, are highlighted with black boxes. **B)** Detailed view of Interface #1, between protomer 1 (light blue) and protomer 2 (green). Residues forming key interactions are depicted as atom-colored sticks and hydrogen bonds depicted as dashed lines. **C)** Detailed view of Interface #1, between protomer 1 (light blue) and protomer 2 (green). Residues forming key interactions are depicted as atom-colored sticks and hydrogen bonds depicted as dashed lines. **D)** Detailed view of Interface #2 between protomer 1 (light blue) and protomer 4 (magenta). Residues forming key interactions are depicted as atom-colored sticks and hydrogen bonds depicted as dashed lines.

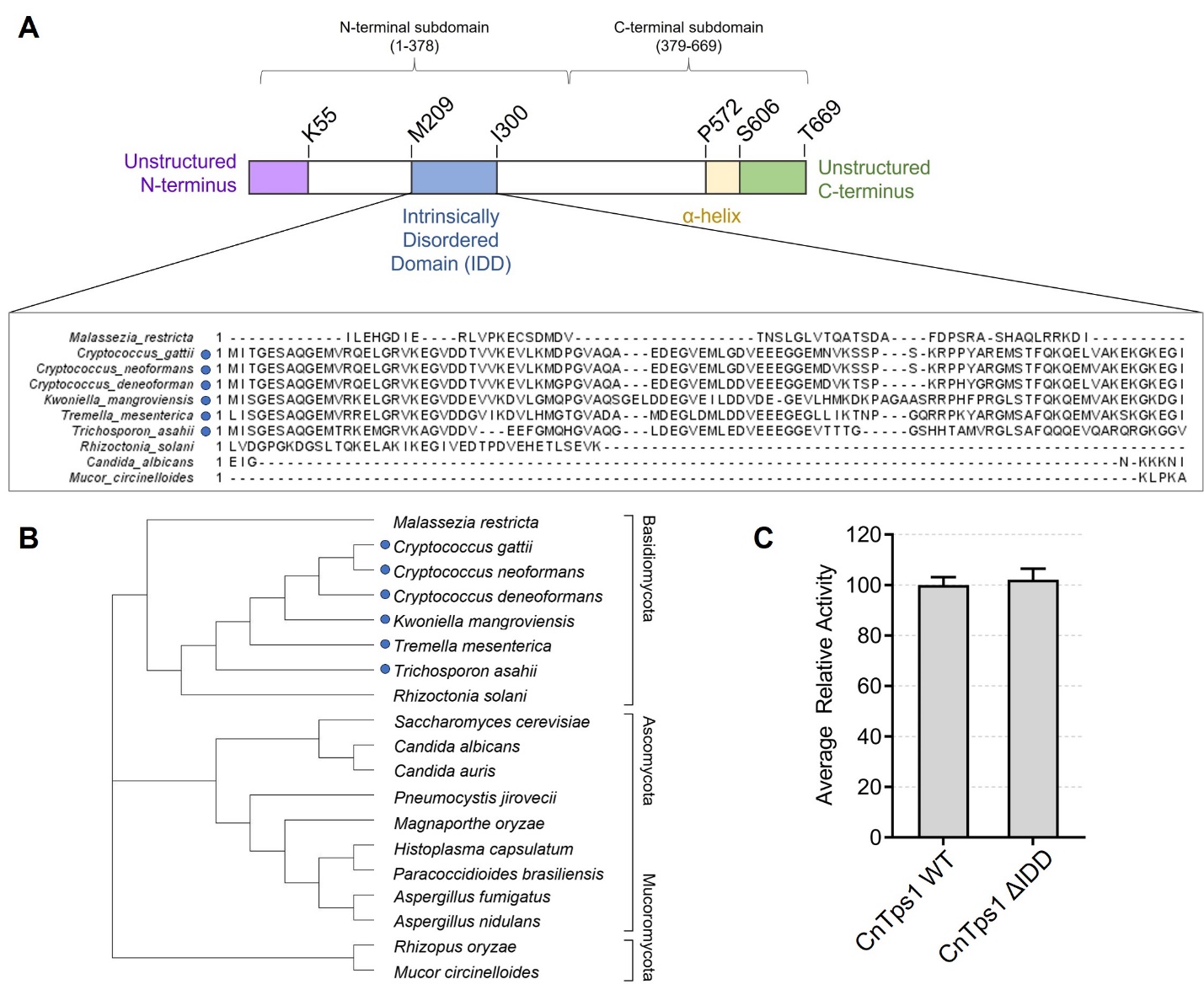

**Figure S13. Characterization of the CnTps1 Intrinsically Disordered Domain. A)** A cartoon of the key features of the *Cryptococcus neoformans* var. *grubii* strain H99 Tps1, detailing the unstructured N and C-termini (purple, green, respectively) and intrinsically disordered domain (IDD, blue) and the α-helix that spans the N and C-terminal subdomains of the protein (yellow). Domains are labelled numerically according to the *C. neoformans* var. *grubii* strain H99 Tps1 amino acid sequence. An alignment of the sequences of the *C. neoformans* var. *grubii* strain H99 Tps1 Intrinsically Disordered Domain (IDD), residues M209 to I300, with other fungal species. Blue circles represent sequences with greater than 75% sequence identity compared to the *C. neoformans* var. *grubii* strain H99 Tps1 IDD. **B)** An unrooted phylogenetic tree of selected fungal species including representative Basidiomycota, Ascomycota and Mucoromycota, based on *C. neoformans* var. *grubii* strain H99 Tps1. Sequences contained greater than 75% sequence identity compared to the *C. neoformans* sequence are marked with a blue circle, corresponding to the alignment in panel A. **C)** Relative enzymatic activity of wild-type CnTps1 and the CnTps1 ΔIDD protein *in vitro*. Error bars represent the standard error of three independent measurements.

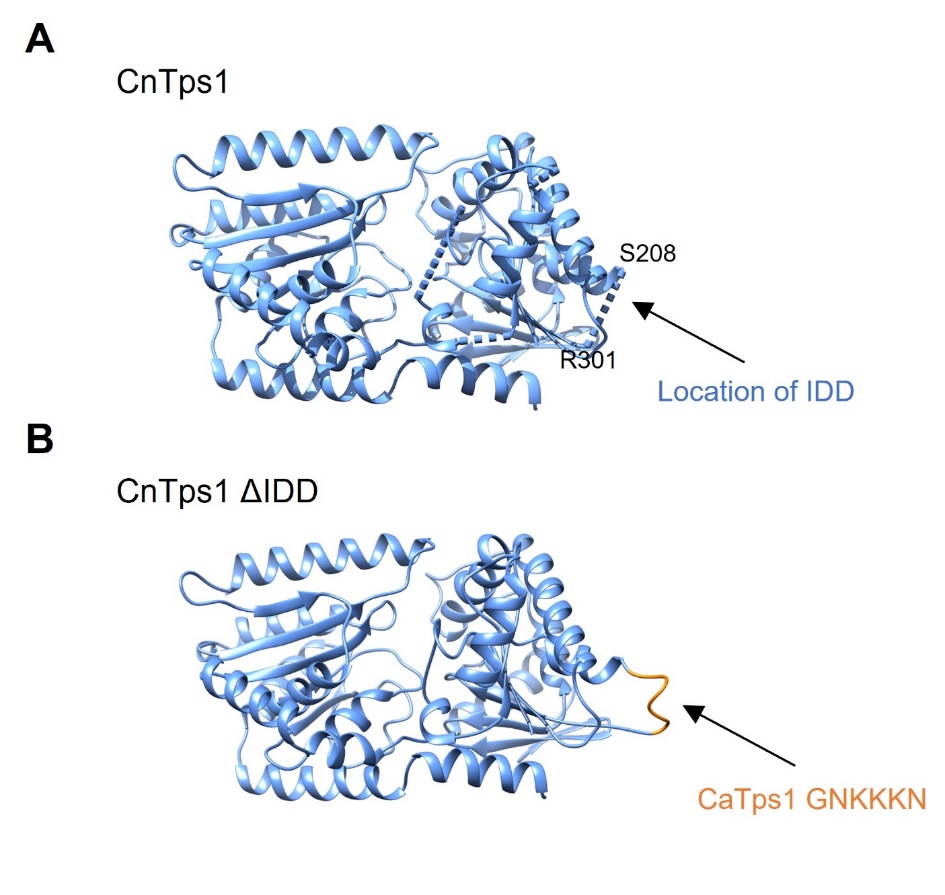

**Figure S14. Generation of CnTps1 ΔIDD protein.**

**A)** Ribbon diagram of CnTps1 protomer in light blue. For clarity, the UDP and G6P molecules are not shown. Dashed lines represent the IDD, M209 – I300, which are not observed in our cryo-EM density maps. **B)** Ribbon diagram of the CnTps1 ΔIDD model, in which the CaTps1 residues (PDB ID 5HUT) in orange (GNKKKN) replace the CnTps1 IDD residues.

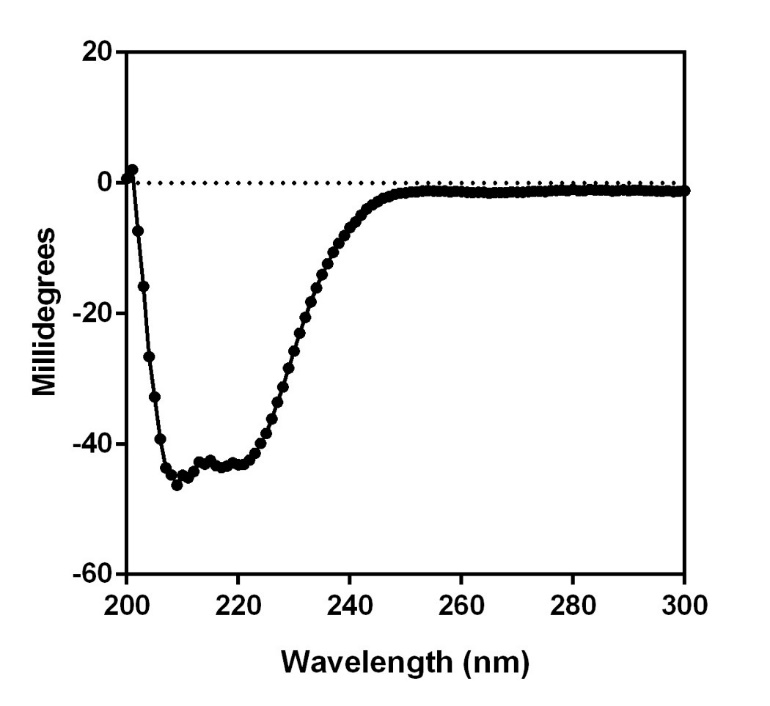

**Figure S15. Circular dichroism (CD) spectrum of CnTps1 ΔIDD.** Circular dichroism spectrum of CnTps1 ΔIDD demonstrates that the mutant is folded.

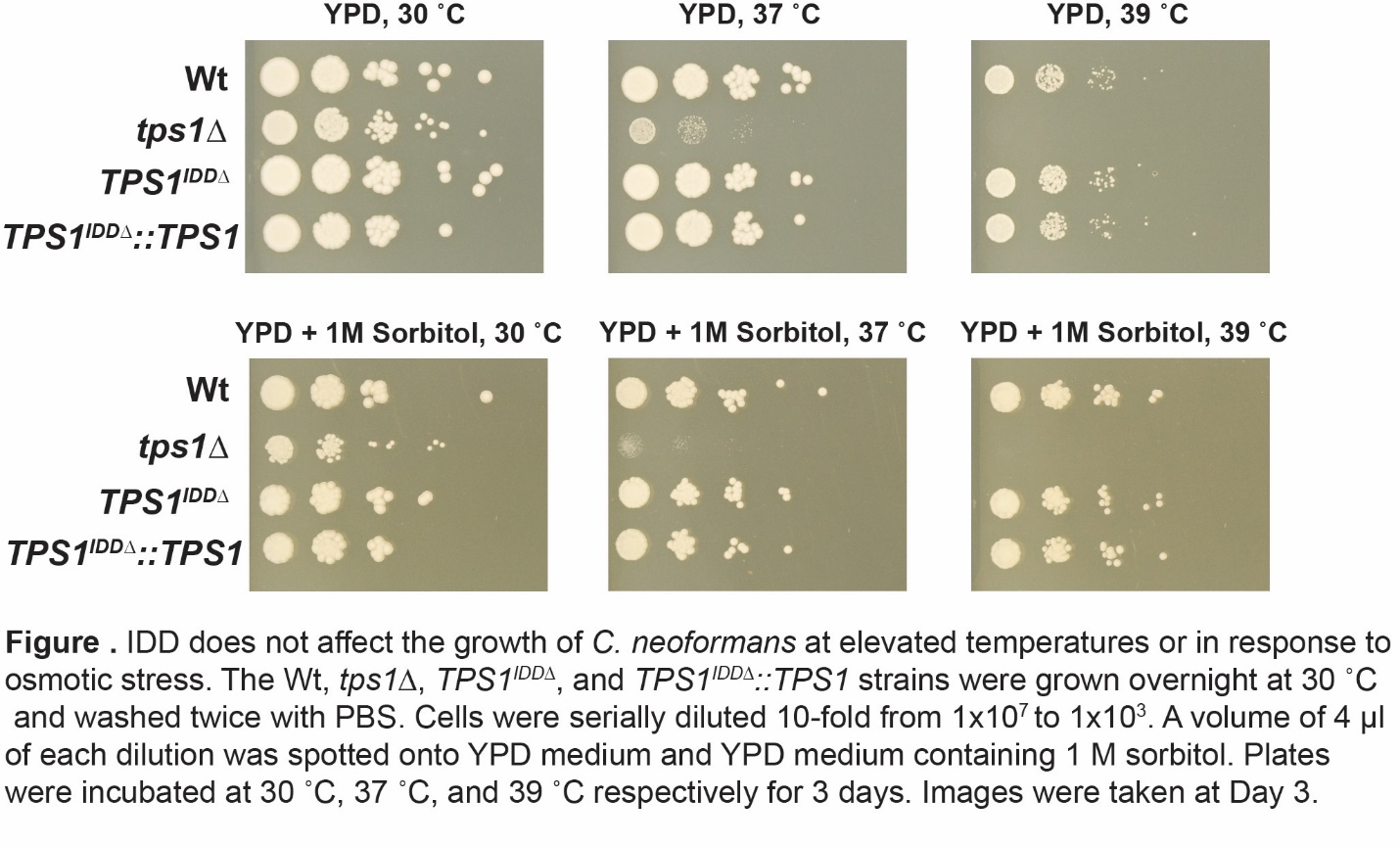

**Figure S16. The CnTps1 IDD does not affect the growth of *C. neoformans* at elevated temperatures or in response to osmotic stress.** The Wt (wild-type), *tps1*Δ, *TPS1^IDDΔ^* and TPS1*^IDDΔ^::TPS1* strains were grown overnight at 30 °C and washed twice with PBS. Cells were serially diluted 10-fold from 1x10^7^ to 1x10^3^. A volume of 4 µL of each dilution was spotted onto YPD medium and YPD medium containing 1 M sorbitol. Plates were incubated at 30 °C, 37 °C and 39 °C respectively for 3 days. Images were taken at Day 3.

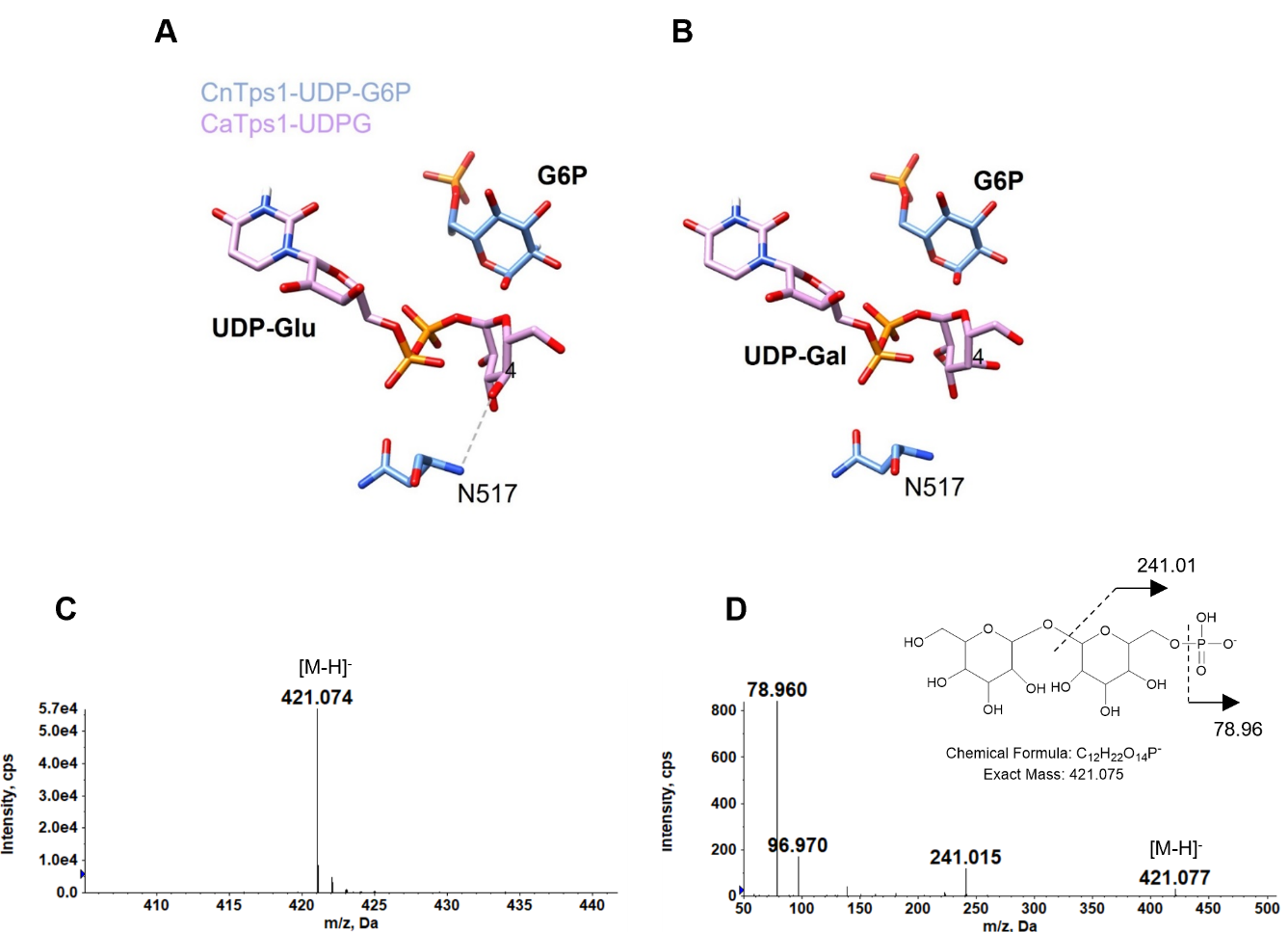

**Figure S17. Details regarding the enzyme specificity of CnTps1.**

**A)** An overlay of CaTps1-UDPG (PDB ID 5HUT) and CnTps1-UDP-G6P with the UDP-Glu shown in lavender and CnTps1-UDP-G6P N517 and G6P depicted as light blue. The O4 hydroxyl on the glucose ring of UDP-Glu is labelled. A dashed line represents the hydrogen bond interaction between CnTps1 N517 and UDP-Glucose. **B)** An overlay of CaTps1-UDPG (PDB ID 5HUT) and CnTps1-UDP-G6P with a model of UDP-Galactose (UDP-Gal) shown in lavender and CnTps1-UDP-G6P N517 and G6P depicted in light blue. The O4 hydroxyl on the glucose ring of UDP-Gal is labelled and cannot engage in a hydrogen bond with N517. **C)** Negative ion ESI mass spectrum of disaccharide–P (in UDPG 4min) **D)** MS/MS of disaccharide–P [M-H]^-^ at *m/z* 421.

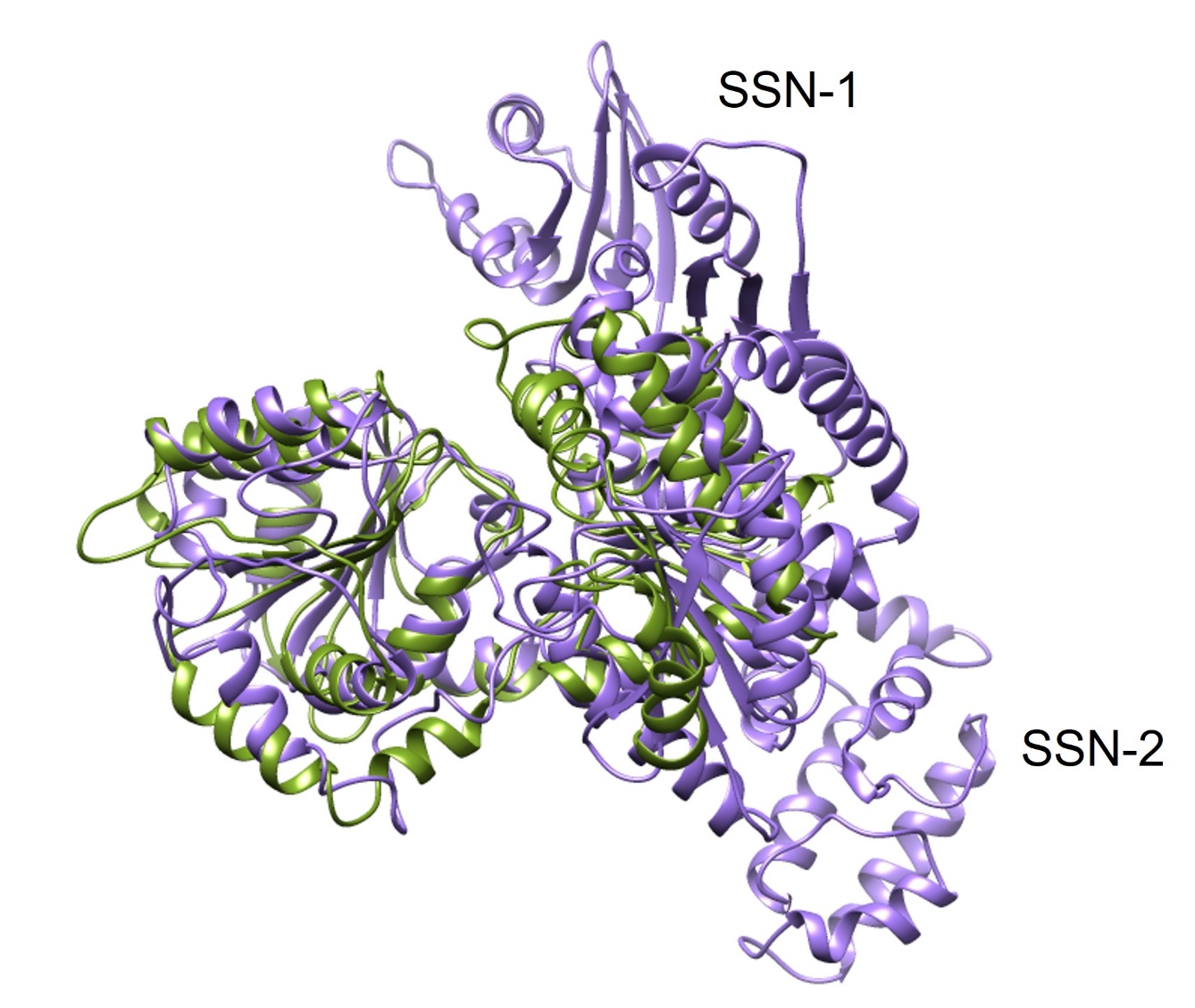

**Figure S18. Comparison of unliganded CnTps1 with sucrose synthase from *Nitrosomonas europaea*.** **A)** An overlay of an unliganded CnTps1 protomer (in green) from the cryo-EM structure with the protomer of *N. europaea* sucrose synthase (purple; PDB ID 4RBN) (5). Dashed lines represent residues that could not be built into the model.

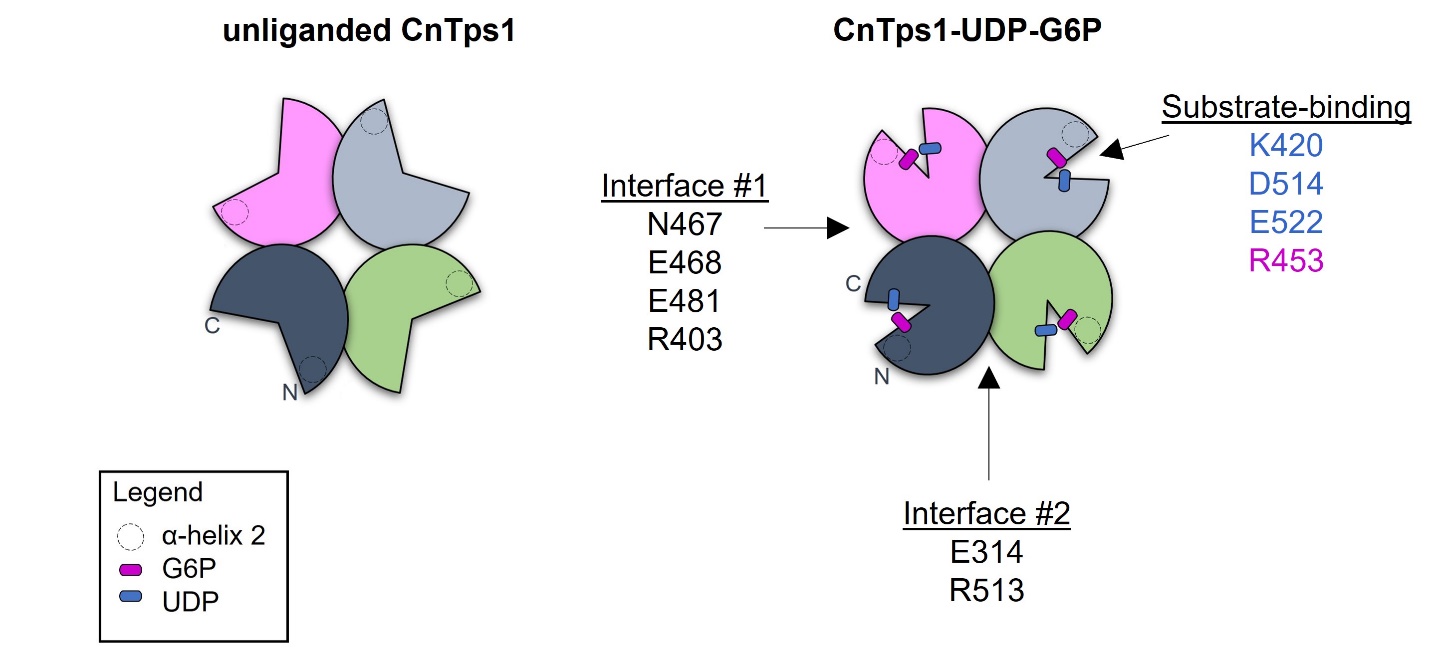

**Figure S19. Proposed model of CnTps1 homo-tetramer function.**

A cartoon depiction of the CnTps1 homo-tetramer in the unliganded and UDP and G6P bound forms is shown. Residues that are involved in tetramerization in CnTps1-UDP-G6P, as well as substrate-binding are highlighted. The locations of the N and C-termini have been shown in a single protomer of each complex. PDBePISA only identified hydrogen bonds between E314 and R513 in interface #2 in *apo* CnTps1.

**Table S1. Michaelis-Menten kinetic parameters of CnTps1 variants and substrates**

| Protein | Substrate | V_max_ (µM/s) | K_m_ (µM) | k_cat_ (s^-1^) | k_cat_/K_m_ (s^-1^ µM^-1^) |
| --- | --- | --- | --- | --- | --- |
| CnTps1 wild-type | UDPG | 14.8 ± 0.8 | 2.6 ± 0.8 | 12.3 | 4.7 |
| CnTps1 ΔIDD | UDPG | 35.5 ± 1.4 | 6.1 ± 1.0 | 29.6 | 4.9 |
| CnTps1 wild-type | UDP-Gal | 6.8 ± 1.0 | 3.9 ± 2.9 | 5.7 | 1.5 |

± represents standard error.

**Table S2. Strains and primers used in this study.**

| **Strains used in this study** | **Background** | **Genotype** | **Reference** |
| --- | --- | --- | --- |
| Wt (H99) |  | *C. neoformans* var. *grubii* MATa | (6) |
| *tps1∆* | Wt (H99) | *tps1∆::NAT* | This study |
| *TPS1^IDD∆^* | Wt (H99) | *NEO-TPS1^IDD∆^* | This study |
| *TPS1^IDD∆^::TPS1* | *TPS1^IDD∆^* | *NAT-TPS1^IDD∆^::TPS1* | This study |
| ***tps1∆*** |  |  |  |
| Primer | 5'-3' sequence | Purpose |  |
| AD131 | TCGGTACCCGGGGATC CAAGCGTACTACTGCAATTAG | to amplify the 5' flank |  |
| AD132 | CATAGCTGTTTCCTG GATCGTTGTTTTTTCTATTTG |  |  |
| AD133 | GAAAAAACAACGATC CAGGAAACAGCTATGACCATG | to amplify the NAT drug selection cassette | |
| AD134 | ACTCATGAATGAAGC GTAAAACGACGGCCAGTGA |  |  |
| AD135 | TGGCCGTCGTTTTAC GCTTCATTCATGAGTACTGG | to amplify the 3' flank |  |
| AD136 | GGTCGACTCTAGAGGATC CTCCCAATAGGCTCCACGC |  |  |
| AD161 | CACTACGTCGTCGTCTGG | to screen for presence/absence of gene | |
| AD162 | ACTGTCAGCAAGCTCATC |  |  |
| AD3858 | TAACGAAGTGCGGCTACCAC | to amplify the *TPS1* locus |  |
| AD3859 | CTGTTAGCGACAATCATTAG |  |  |
| ***TPS1^IDD∆^*** |  |  |  |
| Primer | 5'-3' sequence* | Purpose |  |
| AD3710 | GAGCTCGGTACCCGGGGATC TACGTGCATTTATGTTCAGCCAC | to amplify the 5' flank |  |
| AD3711 | GGTCATAGCTGTTTCCTG GAGATCCTAGGGGCGAATGA |  |  |
| AD3712 | GTCATTCGCCCCTAGGATCTC CAGGAAACAGCTATGACC | to amplify the NEO drug selection cassette | |
| AD3713 | GTTACCGTACGTACTACCTGTTG GTAAAACGACGGCCAGTG |  |  |
| AD3714 | CACTGGCCGTCGTTTTAC CAACAGGTAGTACGTACGGTAAC | to amplify 5' flank and part of *TPS1* gene | |
| AD3715 | GTTCTTCTTCTTGTTACCGAT CATAGAACGGAGCAACATG | and introduce Ca loop sequence |  |
| AD3716 | ATCGGTAACAAGAAGAAGAAC ATCAGGATTGGTTTCTTT | to amplify part of *TPS1* gene and 3' flank | |
| AD3717 | CAGGTCGACTCTAGAGGATC CATCGGCCGTAGATTGGACGTCC | and introduce Ca loop sequence |  |
| AD3720 | GACGCTGCCCACTGGCTCGC | to amplify the IDD region/sequence *TPS1* gene | |
| AD3721 | CCTACCCAATAAGATCACAC |  |  |
| AD3718 | GCGATGTCAAGATACACTGCAGC | to amplify the *TPS1* locus |  |
| AD1066 | CTTTCATGCATCGCCAGAC |  |  |
| AD3718 | GCGATGTCAAGATACACTGCAGC | 5' flank |  |
| AD3713 | GTTACCGTACGTACTACCTGTTG GTAAAACGACGGCCAGTG |  |  |
| AD3712 | GTCATTCGCCCCTAGGATCTC CAGGAAACAGCTATGACC | 3' flank |  |
| AD3719 | CGTGAGAGCTTGATTGATCG |  |  |
| ***TPS1^IDD∆^::TPS1*** |  |  |  |
| Primer | 5'-3' sequence | Purpose |  |
| AD3748 | GAGCTCGGTACCCGGGGATC TACGTGCATTTATGTTCAGCCA | Tps1 5' flank |  |
| AD3749 | GGTCATAGCTGTTTCCTG GAGATCCTAGGGGCGAATGACC |  |  |
| AD3750 | GGTCATTCGCCCCTAGGATCTC CAGGAAACAGCTATGACC | NAT drug resistance cassette |  |
| AD3751 | GTTACCGTACGTACTACCTGTTG GTAAAACGACGGCCAGTG |  |  |
| AD3752 | CACTGGCCGTCGTTTTAC CAACAGGTAGTACGTACGGTAAC | Tps1 5'flank and partial TPS1 gene |  |
| AD3753 | CAGGTCGACTCTAGAGGATC CATCGGCCGTAGATTGGAC |  |  |
| AD3720 | GACGCTGCCCACTGGCTCGC | to amplify the IDD region |  |
| AD3721 | CCTACCCAATAAGATCACAC |  |  |
| AD3718 | GCGATGTCAAGATACACTGCAGC | to amplify the *TPS1* locus |  |
| AD1066 | CTTTCATGCATCGCCAGAC |  |  |
| AD3718 | GCGATGTCAAGATACACTGCAGC | 5' flank |  |
| AD3751 | GTTACCGTACGTACTACCTGTTG GTAAAACGACGGCCAGTG |  |  |
| AD3712 | GTCATTCGCCCCTAGGATCTC CAGGAAACAGCTATGACC | 3' flank |  |
| AD3719 | CGTGAGAGCTTGATTGATCG |  |  |
| AD1066 | CTTTCATGCATCGCCAGAC | for sequencing of *TPS1* locus |  |
| AD2102 | GGATACGCTAGCGAGGTCGAG | for sequencing of *TPS1* locus |  |
| AD2878 | CGCTGGAATTTGGCACCAC | for sequencing of *TPS1* locus |  |
| AD2879 | CGCTCGTGTAGATGTCTTC | for sequencing of *TPS1* locus |  |
| AD2882 | GAGTGGATCGGCAAAGTTG | for sequencing of *TPS1* locus |  |
| AD2883 | ATCTATGATCCTCTCAGAG | for sequencing of *TPS1* locus |  |
| AD2884 | AAGTCTGGAAGTTTGAGCAG | for sequencing of *TPS1* locus |  |
| AD2885 | GCCGTCTCTCGGAGATACAC | for sequencing of *TPS1* locus |  |
| AD3718 | GCGATGTCAAGATACACTGCAGC | for sequencing of *TPS1* locus |  |
| AD3721 | CCTACCCAATAAGATCACAC | for sequencing of *TPS1* locus |  |
| AD3869 | CAACAGGTAGTACGTACGGTAAC | for sequencing of *TPS1* locus |  |
| AD3882 | AAGGGATCAGGATTGGTTTC | for sequencing of *TPS1* locus |  |
| AD3893 | GGTCTCGAGACTCAGCCGAAC | for sequencing of *TPS1* locus |  |
